# Inhibiting Glycan Degradation Prevents HIV-Induced Inflammaging and Cognitive Impairment

**DOI:** 10.1101/2025.11.17.688839

**Authors:** Leila B. Giron, Alejandra Borjabad, Eran Hadas, Janeway Granche, Erika G. Marques de Menezes, Thomas A. Premeaux, Hongxia He, Stephen T. Yeung, Shalini Singh, Courtney Friday, Joshua Glover, Eric Balboa, Derrick Dopkin, Michelle Burrows, Anthony Secreto, Nicolas Skuli, Hiroaki Tateno, Paul W Denton, Frank Palella, Michael J. Corley, Lishomwa C. Ndhlovu, Philip J. Norris, Katherine Tassiopoulos, David J. Volsky, Mohamed Abdel-Mohsen

**Author notes:** Co-first authors (Leila B. Giron, Alejandra Borjabad). Co-corresponding authors: David Volsky, Ph.D. Professor. Icahn School of Medicine at Mount Sinai; New York, NY 10029. Mohamed Abdel-Mohsen, Ph.D. Associate Professor, Northwestern University; Chicago, IL 60611.

## Abstract

Cognitive impairment is a frequent outcome of chronic viral infections linked to premature aging, including HIV. The mechanisms underlying this decline remain poorly understood. Here, we identify pro-inflammatory glycan degradation, characterized by loss of sialic acid and galactose, alterations that are hallmarks of premature aging, as key contributors to HIV-associated cognitive impairment (HIV-CI). In two independent cohorts of people living with HIV, these degradative changes were enriched in individuals with cognitive impairment, particularly females, and correlated with worse cognitive performance. In both a humanized mouse model of HIV and Eco-HIV, a complementary model that allows behavioral testing, pharmacological inhibition of glycan degradation with sialidase inhibitors prevented virally induced inflammation, immune activation, accelerated aging, and memory deficits. These findings implicate glycan degradation as a contributor to inflammation and cognitive impairment in HIV and highlight glycan-preserving therapies as a promising strategy to mitigate inflammation, premature aging, and cognitive decline during viral infections.

## INTRODUCTION

Cognitive impairment is a well-recognized consequence of chronic viral infections, reflecting the combined effects of persistent inflammation, premature aging, and neurobiological disruption. Among these, HIV-associated cognitive impairment (HIV-CI) is the most prevalent and best studied, affecting at least 24% of people living with HIV (PLWH) despite effective viral suppression with antiretroviral therapy (ART).^1–3^ Although often mild, these impairments substantially diminish quality of life, increase long-term health risks, and worsen with aging in PLWH. The physiologic mechanisms driving virally mediated cognitive impairment, including HIV-CI, remain incompletely defined and are likely multifactorial, underscoring the urgent need for deeper mechanistic insights to inform targeted prevention and therapeutic strategies.

A growing body of evidence implicates myeloid cell activation in the establishment of a pro-inflammatory environment driving HIV neuropathogenesis.^4–10^ Yet, the upstream mechanisms sustaining this myeloid-driven neuroinflammation remain unclear. One emerging candidate is aberrant glycosylation, a fundamental regulator of immune signaling and inflammatory tone. Glycan patterns on circulating and cell-surface glycoproteins dictate their interactions with glycan-binding receptors on immune cells, thereby shaping immune cell activation and effector functions.^11–14^ A notable example is the anti-inflammatory activity of intravenous immunoglobulins (IVIGs), which depends on antibody sialylation (the presence of the glycan, sialic acid).^15–23^ Sialic acid on circulating glycoproteins can trigger anti-inflammatory processes by engaging sialic acid–binding proteins on immune cells, inducing REST-mediated repression of NF-κB signaling thereby protecting against excessive inflammation during viral infections.^24^ Conversely, loss of sialic acid (hypo-sialylation) disrupts these pathways and amplifies inflammatory responses.^24^

Consistent with the pro-inflammatory effects of glycomic alterations, hypo-sialylation and agalactosylation (loss of the sugar galactose, which is required for sialylation) have emerged as hallmarks of accelerated biological aging.^25,26^ Together, these alterations amplify inflammation and contribute to immune dysregulation. In line with this biology, agalactosylation and hypo-sialylation are associated with aging-related illnesses, including cardiovascular disease, cancer, and neurodegenerative conditions such as Alzheimer’s disease and vascular cognitive impairment.^27–34^ Moreover, glycosylation disruptions impair neural transmission and protein stability, further implicating them in neurodegeneration and cognitive decline.^35–38^

In PLWH, ART-suppressed individuals exhibit distinct glycomic alterations including hypo-sialylation and agalactosylation.^39–41^ These traits correlate with heightened systemic inflammation, premature biological aging, and impaired Fc-mediated antiviral immunity.^41^ However, whether such glycomic alterations directly contribute to neuroinflammation and HIV-CI has not been determined.

In this study, we address this critical gap. Using two well-characterized cohorts of PLWH on ART, we show that individuals with HIV-CI, particularly females, exhibit more pronounced glycomic degradation, specifically hypo-sialylation and agalactosylation, than those without impairment. We then tested causality using two complementary animal models: a humanized mouse model of HIV infection and a chimeric viral infection model in immunocompetent mice that permits detailed cognitive assessment. In these systems, pharmacological inhibition of sialidases, enzymes that cleave sialic acids from glycans, preserved sialylation, reduced HIV-driven inflammation and accelerated biological aging, and prevented virally induced cognitive impairment. While sialidase inhibitors are used clinically as antivirals against influenza, their potential application as anti-inflammatory agents has not previously been explored. Together, these findings reveal a previously unrecognized role for glycomic alterations in the pathogenesis of HIV-CI and identify glycan-targeting strategies as promising therapeutic avenues to prevent or ameliorate cognitive impairment in PLWH and potentially other virally mediated cognitive disorders.

## RESULTS

### PLWH on ART with cognitive impairment exhibit more pronounced glycomic alterations, including hypo-sialylation and agalactosylation, compared to PLWH without cognitive impairment in two independent cohorts

We began our investigation by analyzing longitudinal samples collected over an 8-year period from the Advancing Clinical Therapeutics Globally for HIV/AIDS and Other Infections (ACTG) study: HIV Infection, Aging, and Immune Function Long-term Observational cohort (HAILO; A5322). This cohort included 40 PLWH on ART (20 males and 20 females) with or without CI, matched by sex, age, and ethnicity (**Table 1**; **Fig. 1a**). Cognitive function was assessed using a standardized neuropsychological battery, including the Trail Making Tests A and B, the Wechsler Adult Intelligence Scale-Revised Digit Symbol Test, and the Hopkins Verbal Learning Test-Revised. A composite NPZ4 score was calculated as the mean of these test scores, and CI was defined as ≥2 test z-scores ≤ –1 SD or at least one test z-score ≤ –2 SD from normative means (**Fig. 1b**).

**Table 1.**
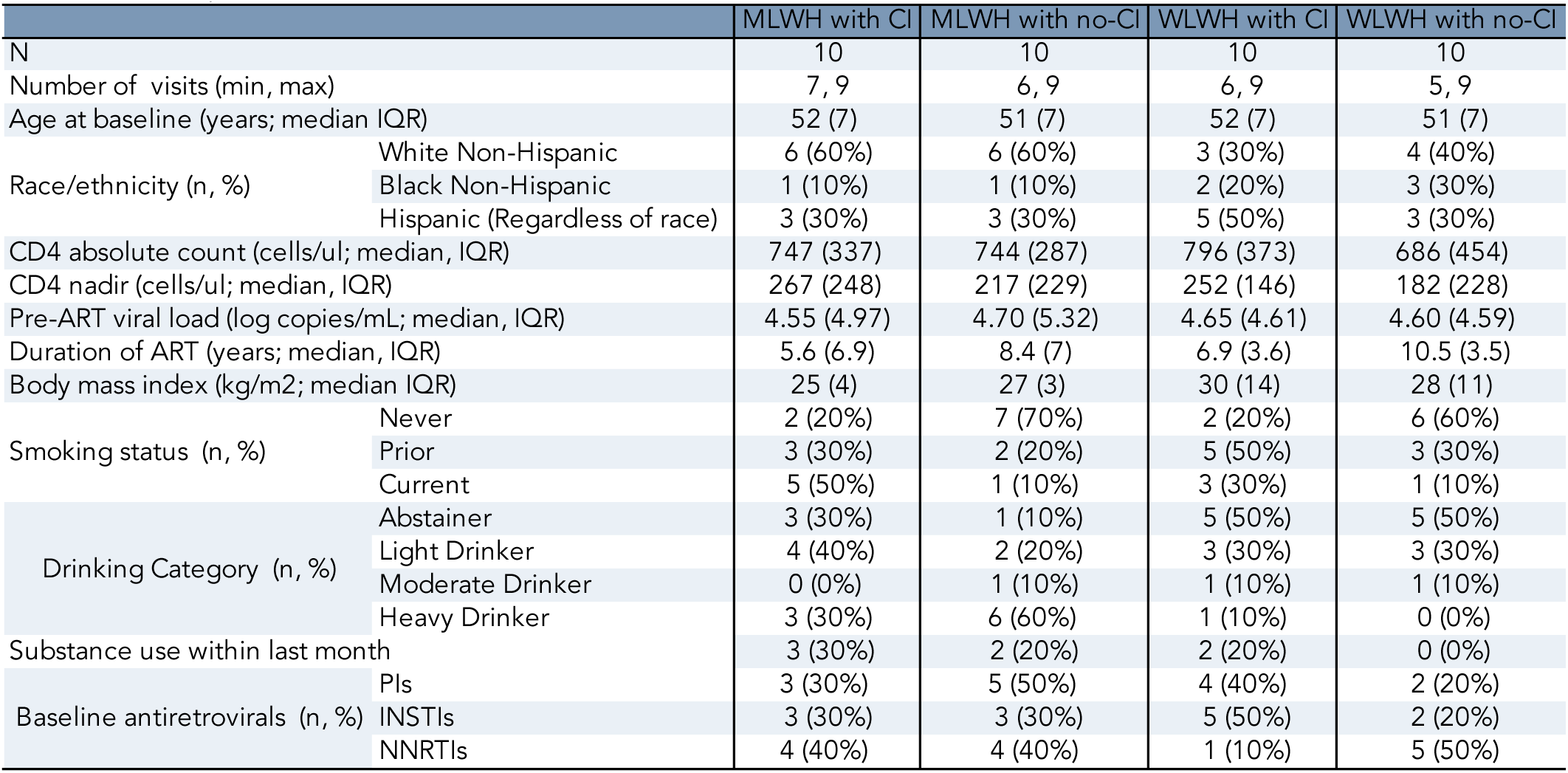
Demographic and clinical characteristics of the testing cohort.

**Figure 1.**
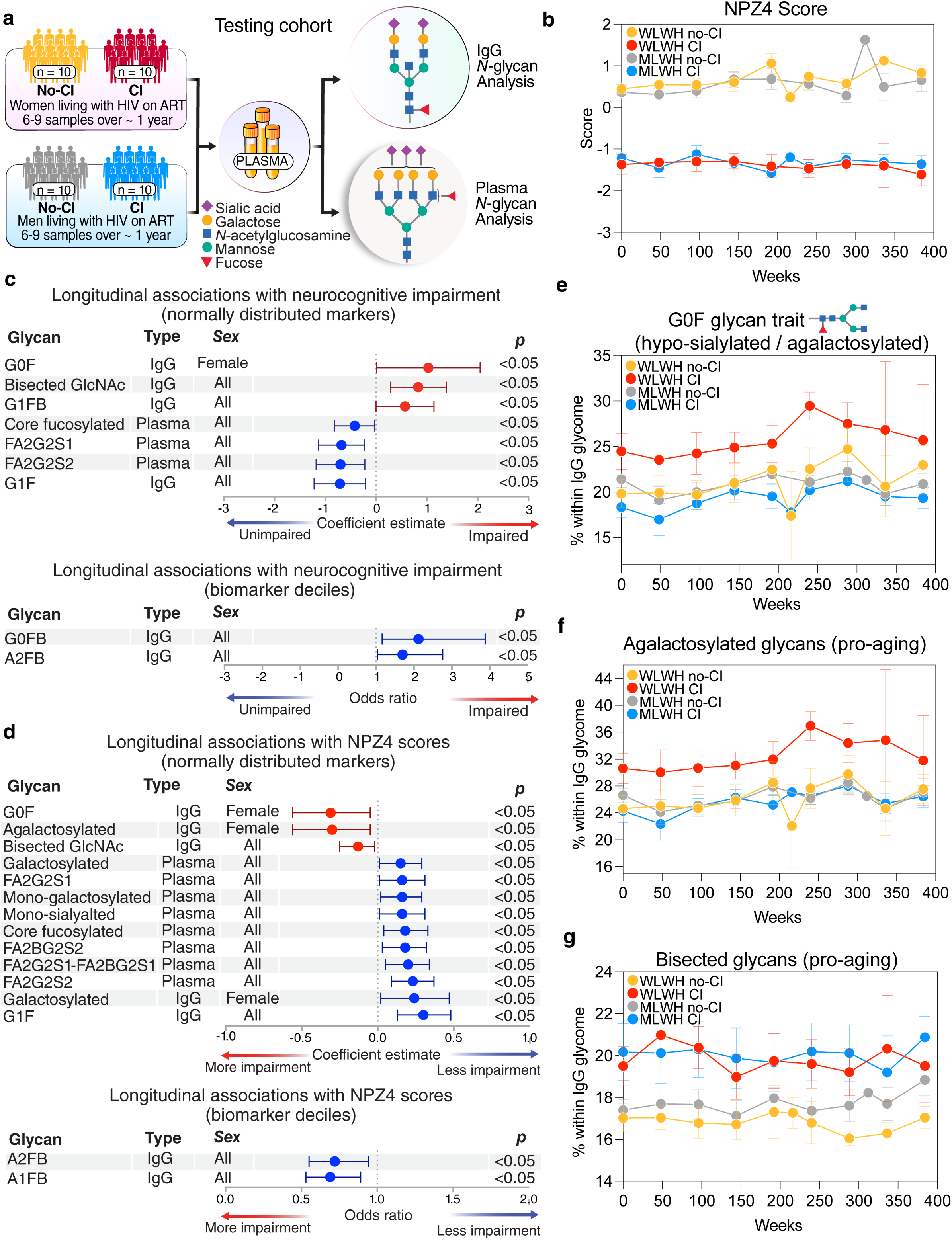
PLWH on ART with cognitive impairment exhibit glycomic alterations, including agalactosylation and hypo-sialylation, compared to PLWH on ART without cognitive impairment in a longitudinal cohort (ACTG A5322 HAILO). **(a)** Schematic of the longitudinal study design, showing IgG and plasma *N*-glycan analyses from 40 females and males living with HIV on ART, with and without cognitive impairment (CI), matched by age, sex, and ethnicity. **(b)** Composite NPZ4 cognitive scores over time. CI was defined as ≥2 test z-scores ≤ –1 SD or ≥1 test z-score ≤ –2 SD. Error bars represent mean ± standard error of the mean (SEM). **(c)** Mixed-effects model results showing longitudinal associations between glycans and CI status. Red = glycans positively associated with CI, blue = glycans associated with no-CI status. **(d)** Mixed-effects model results showing longitudinal associations between glycans and NPZ4 scores. Red = glycans correlated with lower NPZ4 scores, blue = glycans correlated with higher NPZ4 scores. **(e–g)** Representative examples of specific glycan traits from panel (c). Error bars represent mean ± SEM.

Longitudinal plasma samples (5-9 per participant, collected over 8 years) were analyzed for IgG and plasma glycomic profiles using capillary electrophoresis (**Supplementary Fig. 1**). Associations between glycomic traits and cognitive status or NPZ4 scores were assessed using mixed-effects models. Several glycomic alterations were significantly associated with CI or lower NPZ4 scores (**Fig. 1c-g**). For example, pro-inflammatory agalactosylated and hypo-sialylated traits such as G0F, G0FB, and total agalactosylated glycans (which are hypo-sialylated by default) were elevated in PLWH with CI compared to those without impairment, and their levels correlated with reduced NPZ4 scores (indicating poorer cognitive performance). Some of these alterations were more pronounced in women living with HIV (WLWH) than in men (MLWH), although several changes were shared across sexes (**Fig. 1c-g**).

To validate these findings, we next analyzed an independent cross-sectional cohort of 80 PLWH from ACTG HAILO, stratified by sex (40 females and 40 males). Within each sex group, half of the participants had CI and half did not (matched for age and ethnicity; **Fig. 2a**; **Table 2**). In the validation cohort, hypo-sialylated and agalactosylated glycans such as G0F were elevated in individuals with CI compared to those without, while glycans that were galactosylated and sialylated (e.g., A2B) were reduced (**Fig. 2b; Supplementary Fig. 2**). When stratified by sex, these effects were driven primarily by WLWH: WLWH with CI exhibited significantly higher levels of hypo-sialylated and agalactosylated glycans (e.g., G0F and total agalactosylated glycans) and lower levels of galactosylated and sialylated glycans compared to WLWH without CI (**Fig. 2c; Supplementary Fig. 2**). These differences were not observed among MLWH.

**Figure 2.**
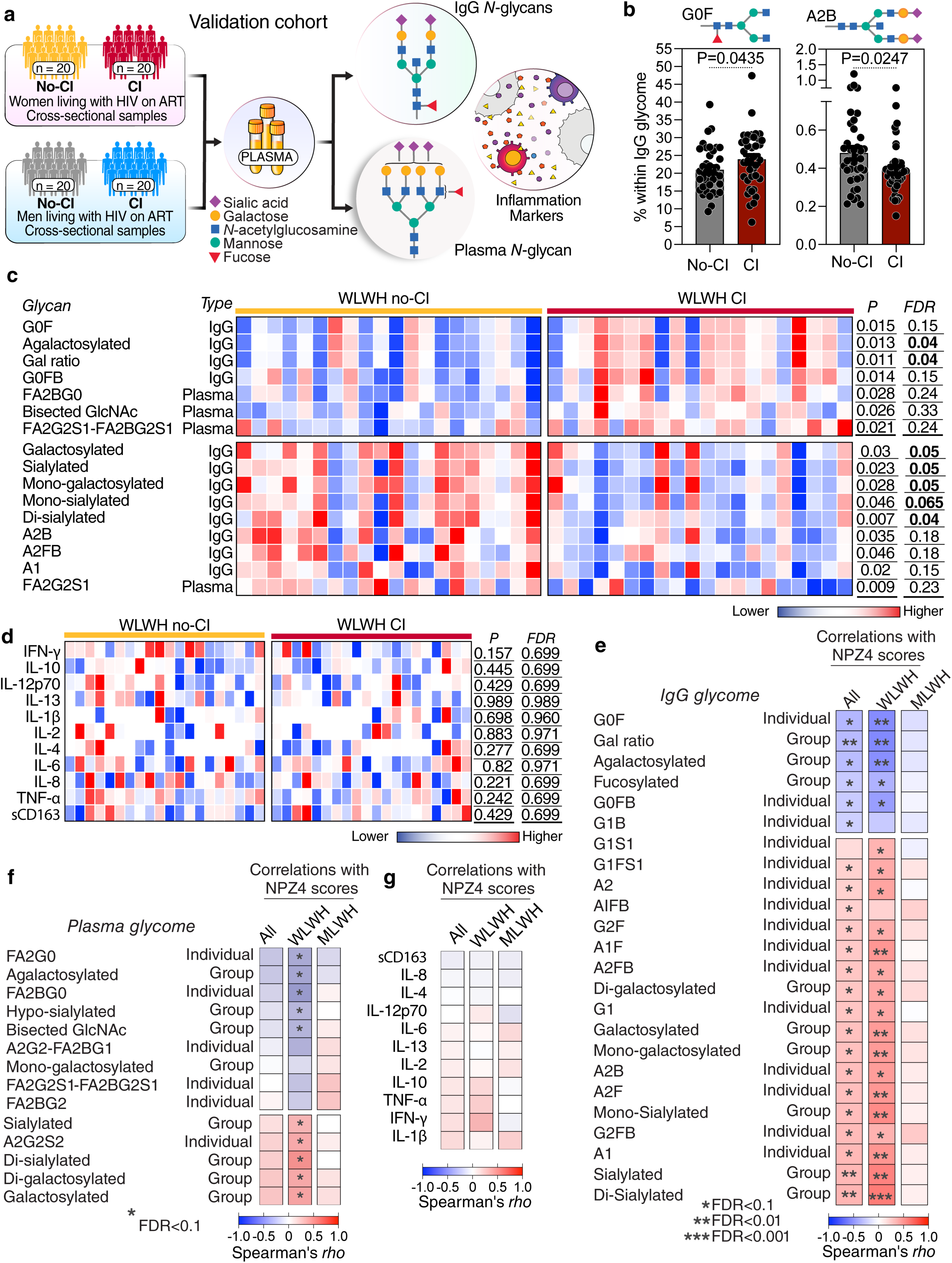
Validation of HIV-CI–associated glycomic alterations in an independent cross-sectional subset of ACTG HAILO. **(a)** Schematic of the validation cohort design (n = 80; 40 females and 40 menmales half with CI and half without), showing both IgG and plasma N-glycan analysis, along with plasma inflammatory marker measurements. **(b)** Representative examples of IgG glycan traits: hypo-sialylated and agalactosylated trait G0F, and sialylated/galactosylated trait A2B, compared between participants with and without CI (males and females combined). Data analyzed using Mann–Whitney tests. Error bars represent mean ± SEM. **(c)** Heatmap of glycan levels in WLWH with and without CI. Red = higher levels, blue = lower levels. Statistical comparisons performed using Mann–Whitney tests with false discovery rate (FDR) correction (Benjamini–Hochberg). **(d)** Heatmap of plasma inflammatory marker levels in WLWH with and without CI. Red = higher levels, blue = lower levels. Statistical comparisons performed as in panel (c). **(e–g)** Spearman’s rank correlation heatmaps of (e) IgG glycan traits, (f) plasma glycan traits, and (g) plasma inflammatory markers with NPZ4 cognitive scores, shown across all participants, WLWH, and MLWH. Rows represent biomarker traits; columns represent correlation with NPZ4 within each subgroup. Red = positive correlation and blue = negative correlation. Significance was assessed with FDR correction (Benjamini–Hochberg).

**Table 2.** Demographic and clinical characteristics of the validation cohort.

|  |  | MLWH with CI | MLWH with no-CI | WLWH with CI | WLWH with no-CI |
| --- | --- | --- | --- | --- | --- |
| N |  | 20 | 20 | 20 | 20 |
| Age (years; median, IQR) |  | 50 (7) | 50 (7) | 51 (12) | 50 (10) |
| Race/ethnicity (n, %) | White Non-Hispanic | 10 (50%) | 10 (50%) | 3 (15%) | 4 (20%) |
|  | Black Non-Hispanic | 4 (20%) | 4 (20%) | 10 (50%) | 12 (60%) |
|  | Hispanic (Regardless of Race) | 6 (30%) | 6 (30%) | 7 (35%) | 4 (20%) |
| CD4 absolute count (cells/μl; median, IQR) | | 740 (369) | 607 (374) | 856 (254) | 771 (417) |
| Nadir CD4 <200 cells/μl (n, %) | | 7 (35%) | 13 (65%) | 8 (40%) | 10 (50%) |
| Pre-ART viral load (log copies/ml; median, IQR) |  | 4.66 (4.94) | 4.65 (5.24) | 4.57 (5.03) | 4.70 (4.96) |
| Duration of ART (years; median, IQR) |  | 8.8 (6) | 11.1 (8.2) | 8.0 (6) | 9.2 (7.9) |
| Body mass index (kg/m<sup>2</sup>) | <18.5 | 0 (0%) | 1 (5%) | 0 (0%) | 0 (0%) |
|  | 18.5 - <25.0 | 5 (25%) | 4 (20%) | 7 (35%) | 3 (15%) |
|  | 25.0 - <30.0 | 6 (30%) | 6 (30%) | 5 (25%) | 3 (15%) |
|  | 30.0+ | 9 (45%) | 9 (45%) | 8 (40%) | 14 (70%) |
| Drinking Category (n, %) | Abstainer | 8 (40%) | 6 (30%) | 8 (40%) | 8 (40%) |
|  | Light Drinker | 7 (35%) | 6 (30%) | 7 (35%) | 8 (40%) |
|  | Moderate Drinker | 0 (0%) | 1 (5%) | 0 (0%) | 2 (10%) |
|  | Heavy Drinker | 5 (25%) | 7 (35%) | 5 (25%) | 2 (10%) |
| Smoking status (n, %) | Never | 9 (45%) | 6 (30%) | 12 (60%) | 8 (40%) |
|  | Prior | 8 (40%) | 10 (50%) | 4 (20%) | 4 (20%) |
|  | Current | 3 (15%) | 4 (20%) | 4 (20%) | 8 (40%) |
| Substance use within last month (excludes alcohol & tobacco) |  | 6 (30%) | 6 (30%) | 4 (20%) | 2 (10%) |
| Baseline antiretrovirals (n, %) | PIs | 2 (10%) | 2 (10%) | 9 (45%) | 7 (35%) |
|  | INSTIs | 9 (45%) | 12 (60%) | 10 (50%) | 10 (50%) |
|  | NNRTIs | 9 (45%) | 7 (35%) | 4 (20%) | 5 (25%) |

Notably, previous research in the general population has identified an IgG *N*-glycan ratio, the Gal-ratio, which reflects the distribution of IgG agalactosylation that can serve as a prognostic biomarker for cancer incidence.^42–44^ Using the same formula, we found that the Gal-ratio was significantly higher in WLWH with CI compared to those without (**Fig. 2c; Supplementary Fig. 2**). To examine the specificity of these associations, we measured 11 inflammatory markers implicated in HIV pathogenesis, including IL-6, TNFα, and IFNγ. Unlike glycomic traits, these inflammatory markers did not differ between WLWH with and without CI (**Fig. 2d**), indicating that glycomic alterations provide a more specific and sensitive biomarker of HIV-associated CI than conventional inflammatory markers.

We next examined for associations between these biomarkers and NPZ4 scores (**Fig. 2e-g**). Agalactosylated and hypo-sialylated traits, as well as the Gal-ratio, were inversely correlated with NPZ4 scores in the overall population, consistent with their pro-inflammatory and detrimental roles. These effects were particularly pronounced among WLWH but not MLWH. In contrast, galactosylated and sialylated glycans correlated positively with NPZ4 scores, reflecting their protective roles. Importantly, inflammatory cytokines did not correlate with NPZ4 scores, further highlighting the distinction between conventional inflammatory markers and glycomic alterations as unique indicators of cognitive function in PLWH.

Together, these findings show that PLWH with CI, particularly females in the cohorts we tested, exhibit distinct glycomic alterations marked by increased degradation of sialic acid (hypo-sialylation) and galactose (agalactosylation), which correlate with worse cognitive performance and provide a more specific signal of HIV-associated CI than conventional inflammatory markers.

### Identification of sialidase inhibitors that prevent glycan loss and reduce myeloid inflammation in vitro

The degradation of key glycans such as sialic acid and galactose, which was increased in PLWH with CI, can result from two processes: reduced expression of glycosyltransferases that add these sugars to glycoproteins and glycolipids, and/or increased expression of glycan-degrading enzymes that remove them. In our recent studies,^41^ glycan loss in PLWH on ART, particularly hypo-sialylation and agalactosylation, was associated with both decreased expression of specific glycosyltransferases and increased levels of glycan-degrading enzymes, including sialidases (which remove sialic acid) and β-galactosidase (which removes galactose). While there are currently no safe pharmacological approaches to inhibit β-galactosidase, sialidase inhibitors have long been used clinically as effective antivirals to block influenza neuraminidase and prevent disease progression.^45,46^ Importantly, these inhibitors also inhibit mammalian sialidases, albeit with lower potency.

Based on this rationale, we investigated whether sialidase inhibitors could reduce HIV-associated inflammation and cognitive impairment. This approach had two objectives: (i) to provide mechanistic evidence that glycan degradation functionally contributes to HIV-associated inflammation and cognitive decline, and (ii) to evaluate the therapeutic potential of repurposing this drug class to prevent HIV-associated inflammation and cognitive impairment.

We first examined oseltamivir (Tamiflu), an FDA-approved influenza neuraminidase inhibitor for its ability to block sialidase-mediated removal of sialic acid from human PBMCs, as measured by binding of the lectin SNA (which specifically recognizes sialic acid). Oseltamivir alone was insufficient, so we tested it in combination with other sialidase inhibitors, including zanamivir (Relenza), peramivir (Rapivab), and the broad-spectrum experimental inhibitor DANA (2,3-dehydro-2-deoxy-N-acetylneuraminic acid). As shown in **Fig. 3a-b**, combinations of these inhibitors effectively prevented sialidase-mediated loss of sialic acid, as indicated by preservation of SNA mean fluorescence intensity (MFI) and SNA-positive cell percentages, without affecting cell viability (**Fig. 3c**).

**Figure 3.**
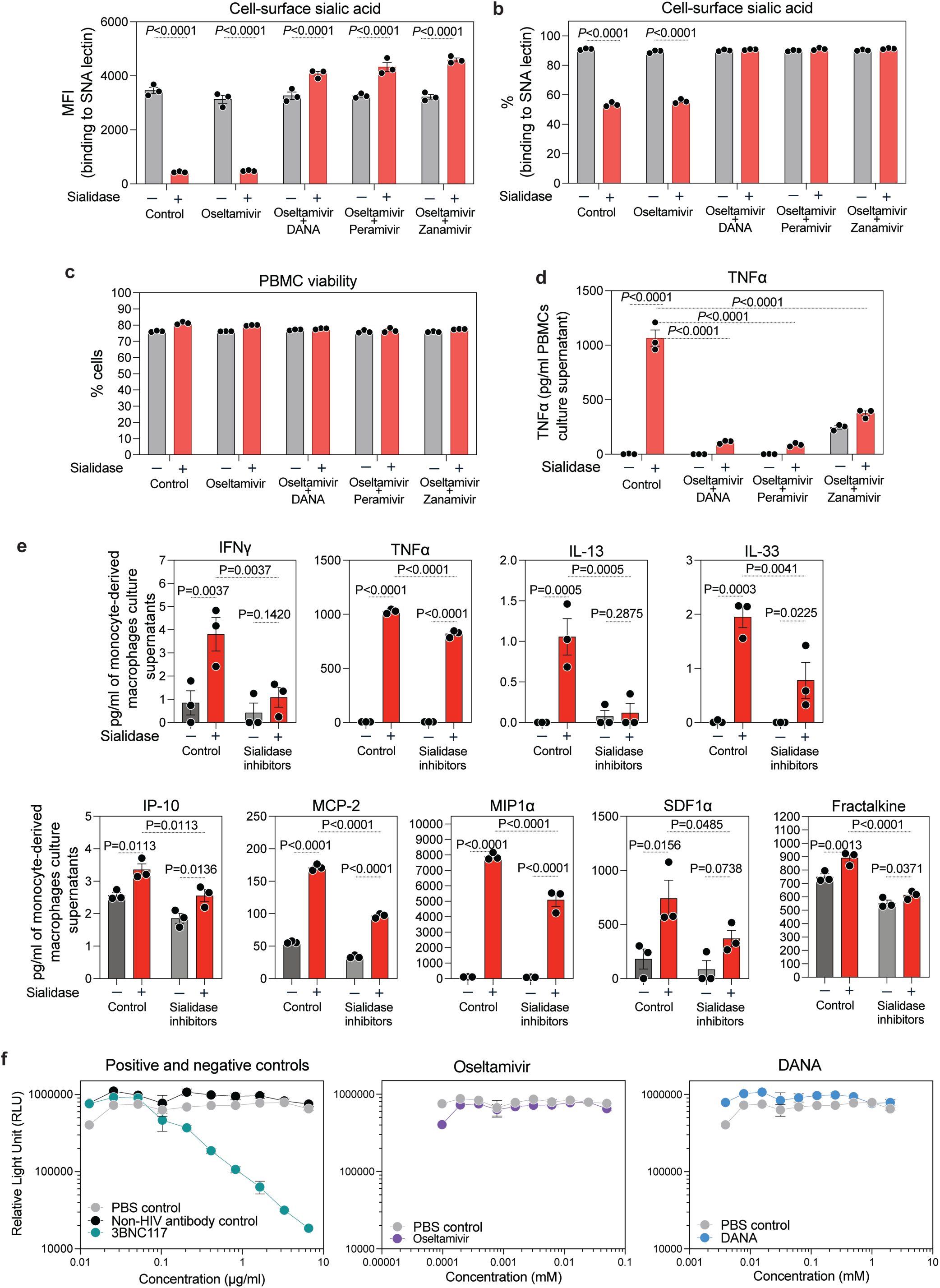
Sialidase inhibitors preserve sialylation and prevent inflammation *in vitro*. **(a–b)** Flow cytometry analysis of sialylation levels on PBMCs treated with exogenous sialidase ± sialidase inhibitors. Sialylation was assessed by SNA lectin binding, measured as mean fluorescence intensity (MFI) (a) and percentage of SNA cells **(b).** Error bars represent mean ± SEM. Statistical analysis by ANOVA with multiple comparisons corrected using the two-stage step-up method of Benjamini, Krieger, and Yekutieli. **(c)** Cell viability of PBMCs under the same conditions, measured by flow cytometry. Error bars represent mean ± SEM. **(d)** Levels of TNFα in culture supernatants of PBMCs treated with sialidase ± inhibitors, measured by ELISA. Statistical analysis by repeated-measures ANOVA with Tukey’s post hoc correction. **(e)** Multiplex cytokine analysis of culture supernatants from monocyte-derived macrophages treated with sialidase in the presence or absence of sialidase inhibitors. Error bars represent mean ± SEM. Statistical analysis by ANOVA with multiple comparisons corrected using the two-stage step-up method of Benjamini, Krieger, and Yekutieli. **(f)** HIV infectivity assays in TZM-bl reporter cells, showing that neither oseltamivir nor DANA affected HIV entry. 3BNC117 antibody was included as a positive control.

As expected, sialidase treatment reduced anti-inflammatory sialylation and increased inflammatory cytokine release, including TNFα, in PBMC supernatants (**Fig. 3d**). In contrast, combinations of sialidase inhibitors prevented this increase in TNFα. We proceeded with the combination of oseltamivir and DANA and tested their effects on monocyte-derived macrophages treated with sialidase. Sialidase exposure induced the expression of multiple inflammatory markers, while co-treatment with sialidase inhibitors prevented sialidase-induced upregulation of inflammatory markers (**Fig. 3e**). These results indicate that sialidase inhibitors, classically used as antiviral agents, also possess previously unrecognized anti-inflammatory properties by preserving sialylation and preventing the degradation of this key anti-inflammatory glycan.

Finally, we tested whether these two inhibitors also had direct antiviral effects on HIV infection using TZM-bl infectivity assays. Neither oseltamivir nor DANA affected HIV infectivity (**Fig. 3f**), compared to PBS (negative control) or the broadly neutralizing antibody 3BNC117 (positive control). Together, these findings suggest that glycan preservation by sialidase inhibitors may represent a novel therapeutic avenue to mitigate HIV-associated inflammation and potentially cognitive impairment.

### Sialidase inhibitors inhibit HIV-mediated inflammation, immune activation, and accelerated biological aging in a humanized mouse model of HIV infection

Having established that combination treatment with sialidase inhibitors is sufficient to prevent cellular glycan degradation *in vitro*, we next examined whether sialidase inhibition with oseltamivir (12.5 mg/kg) and DANA (12.5 mg/kg) could modulate HIV-associated inflammation in vivo using an advanced bone marrow–liver–thymus (BLT) humanized mouse model of HIV infection. Two independent cohorts of BLT female mice, with high levels of human immune cell reconstitution, were infected or not with the HIV_SUMA_ transmitter–founder virus (**Fig. 4a-b**). Mice were treated with a combination of Oseltamivir + DANA or saline control, via daily oral gavage (**Fig. 4a**).

**Figure 4.**
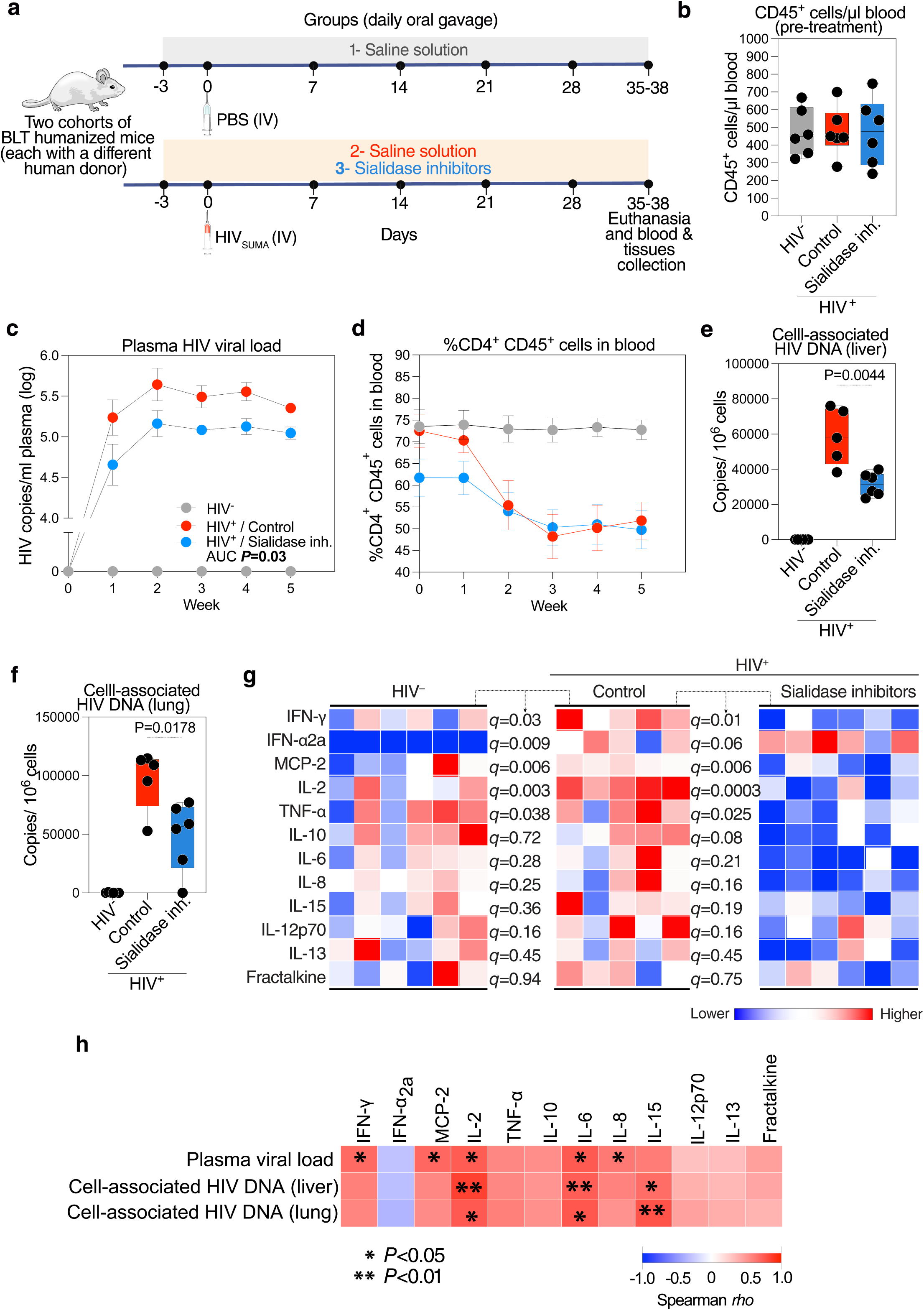
Sialidase inhibition prevents HIV-mediated inflammation in BLT humanized mice. **(a)** Experimental schematic of BLT humanized mouse experiments. Two independent cohorts of BLT mice with robust human immune cell reconstitution were divided into three groups: uninfected controls, HIV-infected untreated, and HIV-infected treated with sialidase inhibitors. Mice were infected with the HIV_SUMA_ transmitter–founder virus. One HIV-infected group was treated daily by oral gavage with Oseltamivir + DANA beginning on day −3 relative to infection, while the other HIV-infected group and the uninfected controls received saline. Blood was collected weekly for five weeks, and tissues were harvested at the study endpoint. **(b)** Human immune reconstitution across groups, shown as % human CD45 cells in blood by flow cytometry. Box-and-whisker plots display all data points. **(c)** Plasma viremia over time. Line graph with mean ± SEM. Treated mice exhibited significantly reduced peak viral loads, analyzed by comparing area under the curve (AUC) between groups using the Mann–Whitney U test (n = 5–6 per group). **(d)** Longitudinal CD4 T cell counts in blood. Line graph with mean ± SEM. Treatment did not prevent HIV-mediated CD4 depletion. **(e–f)** Tissue HIV burden. HIV DNA levels in liver (e) and lung (f), expressed as copies per 10^6^ cells measured by qPCR. Bar graphs are shown as box-and-whisker plots with all points displayed. Unpaired t tests. **(g)** Heatmap of human plasma inflammatory markers measured at study endpoint. Red = higher levels, blue = lower levels. Statistical analysis by ANOVA with multiple comparisons correction. **(h)** Spearman’s rank correlation heatmap showing associations between plasma/tissue viral loads (rows) and plasma inflammatory markers (columns). Red = positive correlation, blue = negative correlation.

Sialidase inhibition reduced peak plasma viremia (**Fig. 4c**) but did not prevent HIV-mediated CD4^+^ T cell depletion (**Fig. 4d**). The treatment also lowered HIV DNA levels in liver and lung tissues compared to controls (**Fig. 4e-f**). Because oseltamivir and DANA do not directly block HIV infectivity (**Fig. 3f**), these effects are likely mediated through anti-inflammatory mechanisms that condition the host environment to be less permissive to viral expansion. Indeed, plasma cytokine profiling at the end of the study showed that, as expected, HIV-infected mice exhibited increased levels of inflammatory markers compared to uninfected mice. Remarkably, treatment with sialidase inhibitors significantly blunted this inflammatory response, as reflected by reduced IFNγ, TNFα, and several other pro-inflammatory cytokines (**Fig. 4g**). Consistent with these findings, plasma viral loads correlated positively with inflammatory markers (**Fig. 4h**), further supporting the link between sialidase inhibition, reduced inflammation, and lower HIV burden.

We next assessed the impact of treatment on markers of T cell activation and exhaustion, hallmarks of HIV pathogenesis. Longitudinal blood and cross-sectional tissue analyses revealed that HIV infection, as expected, increased the expression of activation markers (CD38, HLA-DR, CD69) and exhaustion markers (PD-1) on T cells compared to uninfected controls. Importantly, sialidase inhibitor treatment significantly prevented these HIV-mediated increases in both blood (**Fig. 5a; Supplementary Fig. 3**) and tissues (liver and lung; **Fig. 5b–c**).

**Figure 5.**
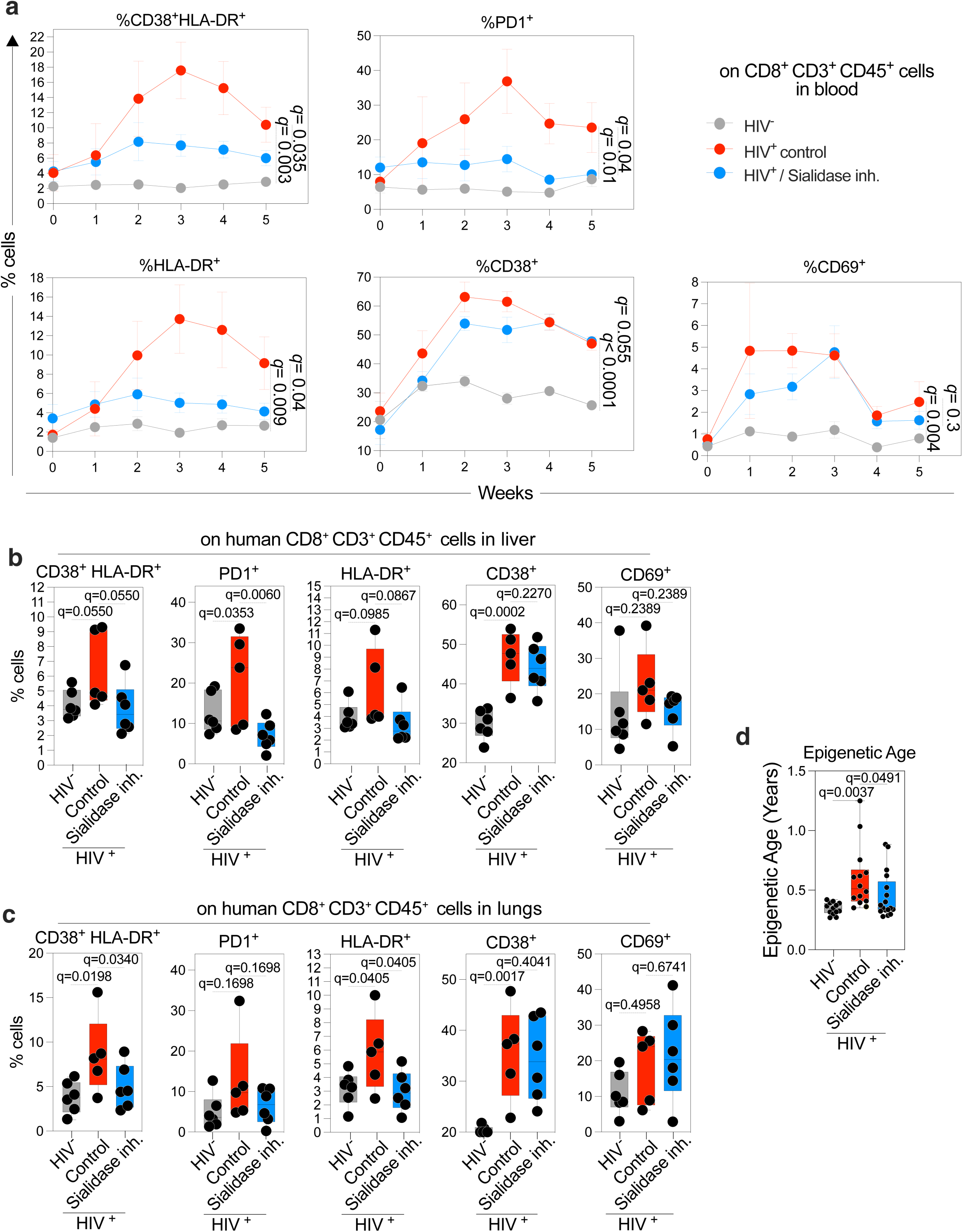
Sialidase inhibition prevents HIV-mediated T cell activation, exhaustion, and acceleration of biological aging. **(a)** Longitudinal line graphs showing the frequency of activated (CD38, HLA-DR, CD69) and exhausted (PD-1) CD8 T cells in peripheral blood of HIV-infected versus uninfected BLT mice, with or without sialidase inhibitor treatment. Treated mice exhibited significantly lower T cell activation and exhaustion over time. Statistical analysis was performed using repeated-measures ANOVA, with area under the curve (AUC) used for multiple comparisons correction. **(b–c)** Box-and-whisker plots showing tissue-resident T cell activation and exhaustion markers in the liver (b) and lungs (c). Sialidase inhibitor treatment significantly reduced immune activation in both tissues. Statistical comparisons performed using ANOVA with multiple comparisons correction by the two-stage step-up method of Benjamini, Krieger, and Yekutieli. **(d)** Boxplots showing epigenetic age acceleration, as measured by DNA methylation clocks, in multiple tissues (lungs, liver, and spleen) from HIV-infected BLT mice. Sialidase inhibitor treatment significantly reduced HIV-associated acceleration in biological aging. Statistical significance was determined by Kruskal–Wallis test.

Because chronic immune activation and inflammation contribute to accelerated biological aging in PLWH, we next examined epigenetic clocks of aging in HIV-infected BLT mice. Sialidase inhibitor treatment significantly reduced HIV-associated acceleration of biological aging in multiple tissues from these mice (**Fig. 5d**). Together, these findings show that sialidase inhibition preserves sialylation *in vivo*, thereby reducing inflammation, immune activation, and epigenetic aging in HIV-infected humanized mice. These results provide mechanistic evidence that glycan degradation contributes directly to HIV-associated pathogenesis, as modulating it produces measurable phenotypic effects, and highlight the therapeutic potential of repurposing sialidase inhibitors to mitigate HIV-associated comorbidities, including cognitive impairment.

### Sialidase inhibitors reduce virally mediated cognitive impairment in the Eco-HIV infected immunocompetent mouse model

While the BLT humanized mouse model provided evidence that sialidase inhibitors can prevent inflammation, immune activation, and accelerated biological aging in the context of a physiologically relevant transmitted-founder strain of HIV using human immune cells, this model is not well-suited to assess neurological or behavioral outcomes. To address this, we employed the model of immunocompetent mice infected with Eco-HIV, a chimeric virus constructed to infect murine lymphocytes, peripheral macrophages, and microglia that has been used to model HIV pathogenesis,^47–58^ inflammation,^59^ neurological impairment,^60–69^ and persistence.^70^ Importantly, in preliminary assessments we found that chronically Eco-HIV infected mice recapitulated key aspects of HIV-associated glycomic dysregulation: brains from these mice had higher levels of NEU1 (encoding sialidase) and lower levels of several sialyltransferases compared to uninfected controls (**Supplementary Fig. 4**), mirroring our previous observations in PLWH and controls.^41^ This finding suggests that the Eco-HIV infected mice are a suitable animal model for testing the role of glycomic alterations in cognitive impairment among PLWH.

We performed three independent experiments with Eco-HIV infected wild-type C57Bl/6j mice. In each, mice were infected or not with Eco-HIV and 25 days after infection, the animals were subjected to daily intranasal treatment with placebo or a combination of oseltamivir (12.5 mg/kg) and DANA (12.5 mg/kg), for 10 days (**Fig. 6a**). Beginning three days after starting drug treatment, mice were tested for cognitive impairment in the radial arm water maze (RAWM) test.^70^ This test measures spatial learning and spatial working memory, two cognitive impairment criteria measured in neuropsychological tests in PLWH,^71^ Across all three experiments, and in combined analysis, treatment with sialidase inhibitors significantly prevented Eco-HIV–mediated spatial learning and spatial memory deficits (**Fig. 6b**).

**Figure 6.**
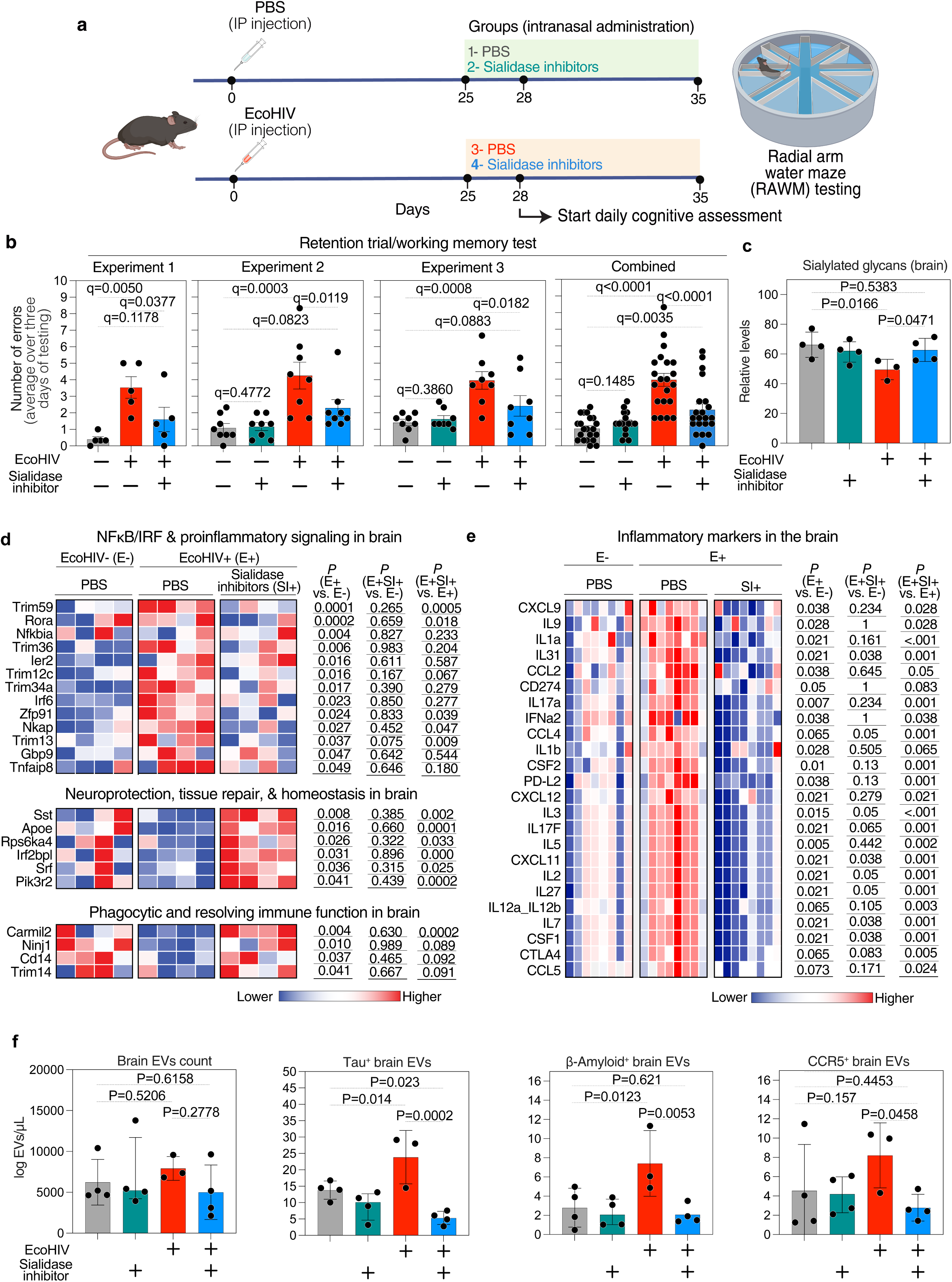
Sialidase inhibitors prevent Eco-HIV–mediated cognitive impairment and neuropathology. **(a)** Experimental schematic of Eco-HIV infection and treatment regimen in C57BL/6 mice. Mice were infected or not with Eco-HIV and, beginning on day 25 post-infection, were treated daily via intranasal administration with a combination of sialidase inhibitors (Oseltamivir + DANA) or PBS. Starting at day 28 post-infection, mice underwent daily cognitive assessments using the radial arm water maze (RAWM) test. Some panels were created with BioRender. **(b)** Bar graphs showing the number of errors in RAWM testing across three independent experiments and combined. Eco-HIV infection significantly impaired learning and memory, while sialidase inhibitor treatment preserved cognitive performance. Data shown as mean ± SEM. Statistical analysis by ANOVA with multiple comparisons correction using the two-stage step-up method of Benjamini, Krieger, and Yekutieli. **(c)** Bar graph showing levels of sialic acid in brain homogenates, measured by SNA lectin binding. Sialidase inhibitor treatment preserved anti-inflammatory sialylation in Eco-HIV–infected mice. Statistical analysis by Fisher’s LSD test. **(d)** RNA-seq heatmap of brain tissues comparing Eco-HIV–infected and control mice, with or without sialidase inhibitor treatment. Eco-HIV infection upregulated NF-κB–related inflammatory pathways and downregulated genes involved in neuroprotection and tissue repair. Treatment normalized these transcriptional alterations. Red = higher expression, blue = lower expression. Unpaired t tests. **(e)** A heatmap of brain tissues showing protein levels of several inflammatory markers in Eco-HIV–infected and control mice, with or without sialidase inhibitor treatment. Eco-HIV infection upregulated several of these markers, while sialidase inhibitors prevented these elevations. Red = higher expression; blue = lower expression. Mann–Whitney T tests. **(f)** Bar graphs showing levels of EVs isolated from mouse brains. Panels include total EVs and EVs expressing neuropathological markers Tau, β-amyloid, and CCR5. Eco-HIV infection increased EV-associated neuropathological markers, which were significantly reduced by sialidase inhibitor treatment. Data shown as mean ± SEM. Fisher’s LSD test.

Neurological improvement was associated with biochemical preservation of anti-inflammatory sialic acid levels in brain tissue from Eco-HIV–infected mice treated with sialidase inhibitors (**Fig. 6c**). Furthermore, RNA-seq analysis of brain tissues revealed that treatment normalized multiple transcriptional changes induced by Eco-HIV infection (**Fig. 6d**). Specifically, Eco-HIV infection upregulated inflammatory gene networks, including NF-κB–related pathways, while downregulating markers of neuroprotection, tissue repair, and immune regulation. Sialidase inhibitor treatment prevented these detrimental transcriptional changes, maintaining reduced expression of inflammatory pathways while restoring neuroprotective, reparative, and immune-related programs. Consistently, analysis of several inflammatory proteins in brain tissues (**Fig. 6e**) and plasma (**Supplementary Fig. 5**) shows that Eco-HIV infection increases the levels of these markers, while sialidase inhibition prevents this Eco-HIV-mediated inflammation. Finally, treatment was associated with reduced levels of neurodegeneration-related markers, including Tau, β-amyloid, and CCR5, measured in extracellular vesicles (EVs) isolated from brain tissue (**Figs. 6f; Supplementary Fig. 6**).

Together, these findings show that sialidase inhibition not only preserves sialylation and reduces neuroinflammation but also protects against Eco-HIV–induced cognitive decline, transcriptional dysregulation, inflammation, and neuropathological markers. These results provide mechanistic evidence linking glycan degradation to HIV-associated cognitive impairment and a preclinical proof-of-concept that repurposed sialidase inhibitors may have therapeutic utility for preventing or treating virally mediated cognitive dysfunction.

## DISCUSSION

In this study, we identified glycan degradation, specifically the loss of the anti-inflammatory sugars sialic acid and galactose, as a potential contributor to inflammation and cognitive impairment in the context of chronic viral infection. In two independent cohorts of PLWH on virally suppressive ART, these glycomic alterations, which are recognized hallmarks of accelerated biological aging,^25,26,41^ were enriched among individuals with cognitive impairment. Notably, these alterations were more pronounced in females, where they correlated strongly with worse cognitive performance. Complementary experiments in two mouse models of HIV infection showed that pharmacological inhibition of glycan degradation with sialidase inhibitors preserved sialylation and prevented systemic and neuroinflammation, prevented accelerated epigenetic aging, and prevented virus-induced memory deficits. Together, these findings bridge human observational data with causal preclinical evidence, implicating glycan degradation as a pathogenic link between viral infection, inflammation, aging, and cognitive decline.

These results highlight glycomic signatures as potential biomarkers of virus-associated cognitive phenotypes. Glycomic alterations were more robustly associated with cognitive impairment than conventional inflammatory cytokines, supporting their sensitivity and specificity as disease indicators. Whether these glycomic features hold prognostic value in predicting the development of cognitive impairment warrants further investigation. Evidence from other contexts supports this possibility: in the general population, glycan degradation has been observed years before the onset of autoimmune conditions such as rheumatoid arthritis,^72,73^ and in PLWH, hypo-sialylation and agalactosylation predicted the development of non-AIDS–defining cancers years prior to diagnosis.^74^ Notably, the IgG “Gal-ratio,” a biomarker of cancer risk,^42–44^ was significantly elevated in females with HIV-associated cognitive impairment. These data raise the possibility that glycomic alterations contribute to a permissive inflammatory state that impairs anti-inflammatory signaling, disrupts immune homeostasis, and increases susceptibility to disease. Longitudinal studies testing whether glycan traits, alone or in combination with inflammatory markers, neuroimaging, or epigenetic clocks, can predict incident cognitive decline will be critical. If validated, glycomic biomarkers could help stratify at-risk individuals before symptoms arise, enabling targeted preventive or therapeutic strategies.

A particularly striking observation was that glycan degradation was most strongly associated with cognitive impairment in females. WLWH who exhibited impairment had significantly higher levels of agalactosylation and hypo-sialylation than unimpaired WLWH, whereas these differences were not consistently observed in males. This sex-specific signal suggests that glycosylation pathways may be differentially regulated by sex hormones, genetic or epigenetic factors, or distinct immune environments. Indeed, estrogen has been shown to directly modulate IgG glycomic profiles, inducing protective anti-inflammatory features such as galactosylation, while menopause accelerates glycan degradation.^41,75–78^ Whether the link between glycomic alterations and cognitive impairment in females is mediated by estrogen activity or other sex-linked mechanisms warrants further study. Importantly, sex differences in glycosylation extend beyond HIV, with implications for autoimmunity, cancer, and neurodegeneration, underscoring the need to better understand how hormonal and genetic factors shape glycan biology in health and disease.

This work also provides insights into the biology of sialic acid and its degradation in viral neuropathogenesis. Sialylated glycans engage sialic acid–binding receptors on immune cells, initiating anti-inflammatory signaling programs and repressing NF-κB activation. Recent findings show that sialylated IgG induces the transcription factor REST in macrophages, thereby repressing NF-κB and protecting against severe influenza disease.^24^ Our results are consistent with this biology: sialidase-mediated loss of sialylation amplified inflammatory pathways and promoted immune activation, whereas pharmacological preservation of sialylation suppressed inflammation and maintained neuroprotective transcriptional programs. These data suggest that the balance between sialylation and sialidase activity is a critical upstream regulator of immune tone during viral infection, and that loss of sialic acid may be a shared mechanism driving chronic inflammation across diverse viral diseases.

The link between glycan degradation and cognitive impairment also intersects with the biology of aging. Hypo-sialylation and agalactosylation are well-established hallmarks of accelerated biological aging, associated with cardiovascular disease, cancer, and neurodegeneration.^27–34^ Our results extend this biology to virally mediated cognitive decline, showing that preservation of sialylation by sialidase inhibition attenuated HIV-associated acceleration of epigenetic aging *in vivo*. These findings support the concept that HIV-associated cognitive impairment may represent, at least in part, a manifestation of “inflammaging,” in which chronic immune dysregulation and glycan degradation accelerate aging processes. By linking viral pathogenesis to conserved aging pathways, our study highlights the potential for glycan-targeting interventions to mitigate both cognitive impairment and broader age-related comorbidities.

Our results further suggest that sialidase inhibitors, widely used as antivirals against influenza, may be repurposed as host-directed therapies that preserve anti-inflammatory glycans. While these drugs do not block HIV infectivity, they maintain host sialylation, thereby suppressing inflammation, immune activation, and biological aging. This shift from direct antiviral to host-directed anti-inflammatory application is novel, and the translational appeal is strengthened by the established safety, clinical use, and accessibility of these agents. Future studies should optimize dosing regimens, test efficacy in chronic infection models, and assess potential synergy with antiretroviral therapy. Beyond HIV, glycan-preserving therapies may have therapeutic value in conditions characterized by chronic inflammation and neurodegeneration.

Several limitations should be acknowledged. While two independent human cohorts were analyzed, larger and longitudinal studies will be required to validate glycomic signatures as predictive biomarkers and to clarify how sex, gender, menopause status, and other demographic or clinical factors modulate these associations. Longitudinal analysis of samples collected before and after cognitive impairment develops will be critical to determine whether glycomic alterations are causally predictive. Although Eco-HIV establishes chronic infection in wild-type mice while preserving immune competence and reproducing many pathobiological features observed in PLWH on ART, and enables validation of HIV-associated cognitive disease,^47–70^ it may not fully recapitulate the complexity of human HIV neuropathogenesis. Finally, while sialidase inhibitors preserved glycans and reduced inflammation *in vivo*, their potency and specificity against mammalian sialidases remain limited. Development of next-generation inhibitors with greater selectivity will be important to enhance translational potential.

In summary, this study establishes glycan degradation as a driver of virally mediated inflammation, accelerated aging, and cognitive impairment. HIV infection provides a powerful exemplar, bridging human cohort data with causal preclinical models, but the implications may extend broadly to other viral infections, chronic inflammatory conditions, and age-related disorders. By identifying sialidase activity and glycan loss as upstream regulators of inflammaging and neurodegeneration, we highlight glycan-preserving therapies as a promising strategy to mitigate the neurological and systemic consequences of chronic viral infections. These findings open new avenues for biomarker development, mechanistic discovery, and therapeutic intervention at the intersection of virology, aging, and neuroimmunology.

## MATERIALS AND METHODS

### Study participants and ethics statement

Samples were obtained from a subset of ACTG HAILO participants. The study protocols were approved by the Institutional Review Board of the participating institutions of the ACTG HAILO study. Written informed consent was obtained from all participants. Recruitment and/or sampling of study participants was not conducted specifically for this study. All human research was performed in accordance with the guidelines of the U.S. Department of Health and Human Services

For the longitudinal cohort, samples from 20 cases (10 males and 10 females) with cognitive impairment at all time points and 20 matched controls who did not have cognitive impairment at any time-point were analyzed. Participants were matched on the number of time points, sex (at birth), age, race, ethnicity, and baseline CD4 count. All participants were on ART for ≥ 1 year at baseline (defined as HAILO entry), had a weight of ≤ 300 pounds, a CD4 count of> 350 cells/μl, and an HIV RNA of <50 copies/ml at all visits. Participants contributed 5-9 time points. For the validation cohort, another independent subset of the HAILO cohort was selected: 20 males with cognitive impairment and 20 females with cognitive impairment were matched with 20 males and 20 females who did not have cognitive impairment. Participants were matched on sex (at birth), age (within age group, and within 3 years), race and ethnicity, and nadir CD4 count (≥200 vs <200 cells/mm^3^). For both cohorts, all participants were on ART for ≥1 year at the sampling date, had a weight of ≤300 pounds, a CD4 count of>350 cells/μl, and an HIV RNA of <50 copies/ml.

Cognitive functioning was assessed using the Trail Making A and B tests, the Wechsler Adult Intelligence Scale-Revised Digit Symbol test, and the Hopkins Verbal Learning Test–Revised. Raw scores from each of the four tests were normalized by age, sex, race/ethnicity (black non-Hispanic, white non-Hispanic, or Hispanic), years of education, and learning effects as Z-scores. The NPZ4 score was calculated as the mean of the four normalized scores. Cognitive impairment was defined as having two or more z-scores that were ≥1 standard deviation (SD) below the mean, or at least one z-score that was ≥2 SD below the mean.

### Glycomic analyses

For IgG glycosylation analysis, IgG was isolated from 50 µl of plasma using the Pierce Protein G Spin Plate (Thermo Fisher; Catalog# 45204) and quantified using BCA assay kit (Thermo Fisher; Catalog# 23227). *N*-glycans were released from IgG using peptide-N-glycosidase F (PNGase F) and labeled with 8-aminopyrene-1,3,6-trisulfonic acid (APTS) using the GlycanAssure APTS Kit (Thermo Fisher, Catalog# A33952), following the manufacturer’s protocol. The labeled *N*-glycans were analyzed using the 3500 Genetic Analyzer capillary electrophoresis system. For total plasma protein glycosylation, 18.3μL of plasma was used. *N*-glycans release and labeling were done using the same kit. The relative abundance of *N*-glycan structures was quantified by calculating the area under the curve of each glycan structure divided by the total glycans using the Applied Biosystems GlycanAssure Data Analysis Software Version 2.0, as recently described in details.^41^

### Measurement of inflammatory markers from the human validation cohort

Plasma levels of IFN-γ, IL-1β, IL-2, IL-4, IL-6, IL-8, IL-10, IL-12p70, IL-13, TNF-α were measured using V-PLEX Proinflammatory Panel 1 Human Kit (Meso Scale Diagnostics Catalog# K15049D-1).

### In vitro experiments and TZM-bl infectivity assays

PBMC from healthy donors were cultured in RPMI media (Corning; Catalog# MT10040CM) supplemented with 10% FBS (Thermo Fisher; Catalog# A5669701) and 1% Pen/Strep/Fungizone (Cytiva HyClone; Catalog# SV3007901). PBMC were seeded at 1.75 million cells/ml and treated with a combination of 0.05mM of Oseltamivir (Sigma; Catalog# SML1606) and 2mM of Zanamivir (Cayman Chemical; Catalog# 15123), 2mM of Peramivir (MedChemExpress; Catalog# HY-17015), or 2mM of DANA (Cayman Chemical; Catalog# 19939) for 30 minutes. After incubation, 5μL of sialidase was added to the wells and incubated for 1 hour at 37°C. After sialidase treatment, the cell supernatant was saved for cytokine analysis. Cells were collected and washed in cold PBS for SNA staining. Cells were incubated for 30 minutes at 4°C with 10ug/ml of SNA lectin (Vector Labs; Catalog# FL-1301-2) in PBS containing calcium and magnesium. After incubation, cells were washed with PBS-2%FBS, fixed in PFA (Fixation buffer, Biolegend; Catalog# 420801), and analyzed by flow cytometry (BD Biosciences LSRII). Cytokines were measured in the supernatant using U-PLEX kits from Meso Scale Diagnostics (Biomarker Group 1 (hu) Assays; Catalog# K151AEM-2, Custom Immuno-Oncology Group 1 (hu) Assays; Catalog# K15067L-2) according to manufacture instructions.

For the infectivity assay, TZM-bl cells (NIH AIDS reagent; Catalog# ARP-8129) were seeded at 150,000 cells/mL in R10 with 15μg/mL of dextran and treated with serial dilutions of Oseltamivir or DANA, as well as 3BNC117 as a control. HIV JRCSF was then added to each well and incubated at 37°C for 48 hours. After incubation, 100μL of the cell culture media was added to 100μL of Bright-Glo luciferase (Promega; Catalog# E2610), and the mixture was incubated for 2 minutes. The luciferase–cell media was mixed, and 150μL was transferred to a clear-bottom, black 96-well plate, which was then read in a luminometer immediately.

### Humanized mouse model experiments

BLT mice were generated by the Stem Cell and Xenograft Core (RRID:SCR_010035) in accordance with the University of Pennsylvania Animal Care and Research Committee regulations (IACUC protocol# 803506). Briefly, 6-8 weeks old female NSG (NOD.Cg-*Prkdc^scid^ Il2rg^tm1Wjl^*/SzJ; Jackson Laboratory) mice were injected via the tail vein with CD34^+^ hematopoietic stem cells isolated from human fetal liver tissues. One week post-transplant, human fetal thymic tissue fragments and fetal liver tissue fragments were implanted under the murine renal capsule. Human fetal liver and thymus tissues were procured from Advanced Bioscience Resources (Alameda, CA). Sixteen weeks post-transplant, human immune cell reconstitution in peripheral blood was determined using a LSRII flow cytometer (BD Biosciences, San Jose, CA). Peripheral blood was stained with a live/dead marker (Fixable Viable Stain 510; BD Biosciences, Catalog# 564406), followed by blocking with CD16/CD32 Fc (BD Biosciences, Catalog# 553142) and staining with antibodies for cell surface markers: hCD3-BV421 (BD Biosciences, Catalog# 562426), hCD19-APC (BD Biosciences, Cat No: 555415), hCD45-PE (BD, Catalog# 555483), mCD45-PE-Cy7 (BD Biosciences, Catalog# 552848) and hCD33-BB-515 (BD Biosciences, Catalog# 564588). CountBright absolute counting beads were used according to manufacturer’s instructions (ThermoFisher, Catalog# C36950) and engraftment levels in the xenotransplanted mice were expressed as number human CD45-labeled cells per ul of blood. Engraftment data were analyzed with FlowJo (FlowJo LLC, Ashland, OR). Humanized BLT mice with high immune reconstitution were randomly divided into three groups and each mouse was inoculated intravenously (IV) with 1×10^4^ TCID_50_ of HIV_SUMA_. Peripheral blood was collected weekly for flow cytometry and plasma viral load assay.

### Viral load assessment and cell-associated HIV DNA in humanized mice

Mice were bled weekly, and the blood was centrifuged at 2000rpm for 10 minutes to separate plasma and cells. Plasma was removed and used for viral load and cytokine measurement, while cells were stained as described below. For the viral load, each plasma sample was spiked with a fixed quantity of replication-competent avian leukosis virus (ALV) long terminal repeat (LTR) with a splice adaptor (RCAS) to serve as an internal control.^79–82^ Viral RNA was extracted using QIAamp Viral RNA Mini kit (Qiagen; Catalog# 52906). From RNA, cDNA was generated using the VILO master mix (Thermo Fisher; Catalog# 11755050). Plasma HIV RNA was then amplified by qPCR using LTR-specific primers F522-43 (5’ GCC TCA ATA AAG CTT GCC TTG A 3’) and R626-43 (5’ GGG CGC CAC TGC TAG AGA 3’) coupled with a FAM-BQ probe (5’ CCA GAG TCA CAC AAC AGA CGG GCA CA 3’). The reaction was carried out in a final volume of 20μL, containing 1x TaqMan Universal Master Mix II, including UNG (Applied Biosystems; Catalog# 4440038), 4pmol of each primer and probe, and 5μL of the sample. Cycling conditions were 50°C for 2 min, 95°C for 10 min, followed by 60 cycles of 95°C for 15s and 59°C for 1 min. RCAS internal control was also amplified with the same conditions using specific primers RCAS forward (5’GTC AAT AGA GAG AGG GAT GGA CAA A 3’), RCAS Reverse (5’ TCC ACA AGT GTA GCA GAG CCC 3’), and RCAS Probe FAM-TAMRA (5’ TGG GTC GGG TGG TCG TGC C 3’). By the end of the experiment, tissues were collected, and a single-cell suspension was generated using tissue dissociation kits for lung (Miltenyi, Catalog# 130-095-927) and liver (Miltenyi; Catalog# 130-105-807) in the gentleMACS Octo Dissociator according to the manufacturer’s instructions. DNA and RNA were extracted using AllPrep DNA/RNA/miRNA Universal Kit (Qiagen; Catalog# 80224). Cell-associated HIV DNA was measured by qPCR using the same primers and conditions described above. Human cell count was also estimated by qPCR using human copy number reference assay TERT (Applied Biosystems; Cat# 4403315).

### Flow cytometry assessment of immune activation in the BLT mice

The mouse’s blood was centrifuged at 2000 rpm for 10 minutes, and the plasma was removed. An equal volume of PBS was then added to the tube. Human and mouse Fc blocker (Biolegend; Catalogs# 422301 and 101319) were added to the blood and incubated for 10 minutes. After blocking, cells were incubated with an antibody cocktail containing CD45-AL700 (clone: 2D1, Biolegend; Catalog# 368513), CD8-FITC (clone: SK1, Biolegend; Catalog# 344703), CD38-APC (clone: HIT2, Biolegend; Catalog# 303509), HLA-DR APC-H7 (clone: G46-6, BD Biosciences; Catalog# 561358), CD4-V450 (clone: RPA-T4, BD Biosciences; Catalog# 560345), PD-1-PE (clone:EH12.2H7, Biolegend; Catalog# 329905), and CD3-PE-CF594 (clone: UCHT1, BD Biosciences; Catalog# 562310), CD25-PerCP-Cy5 (clone:BC96, Biolegend; Catalog# 302625), and CD69-PE-Cy7 (clone:FN50; Biolegend; Catalog# 310911) for 30 minutes at room temperature and protected from light. After incubation, red blood cells were lysed using BD FACS Lysing Solution (BD; Catalog# 349202) for 15 minutes. Cells were then washed twice with PBS with 2% FBS, fixed with PFA (Fixation buffer, Biolegend; Catalog# 420801), and analyzed on BD Biosciences LSRII flow cytometer (gating strategy is in **Supplementary Figure 3**).

### Epigenetic clocks of aging

The mammalian DNA methylation array (Illumina HorvathMammalianMethylChip40 BeadChip) available from the Clock Foundation was utilized to obtain DNA methylation data for calculating epigenetic clock estimates.^83^ This platform captures 37,488 CpGs at conserved sequences. Raw data was normalized using the SeSAME R package.^84^

### Eco-HIV-infected mice experiments

EcoHIV was constructed as described using a plasmid encoding HIV/NDK kindly provided by I. Hirsch (Institut National de la Santé et de la Recherche Médicale, Marseille, France).^54^ EcoHIV and ecotropic MLV virus stocks were prepared as previously described.^70^ All animal studies were conducted with the approval of the Icahn School of Medicine at Mount Sinai Institutional Animal Care and Use Committee (IACUC) in full compliance with the U.S. Animal Welfare Act and Public Health Service (PHS) policies. Adult C57BL/6 mice were purchased from Jackson Laboratory (Bar Harbor, ME). Animals were maintained under standard mouse husbandry conditions. Discomfort, distress, and injury to the animals were minimized. Mice were infected with EcoHIV by intraperitoneal inoculation. EcoHIV, 0.5 ml solution of virus stock was injected into the peritoneal cavity as described.^49^ Control animals received 0.5 ml of PBS. Mice were inoculated at 6 to 8 weeks of age with a dose of 2×10^6^ pg/ml of EcoHIV.

### Radial arm water maze (RAWM)

To test spatial learning and memory abilities, we used a radial arm water maze (RAWM) test as previously described.^63,70^ Briefly, RAWM consisted of a six-arm water maze including a hidden platform beneath opaque water and a set of visual cues at the end of each maze arm. The platform is randomly changed on each day of testing to analyze working memory at the end of the day. The testing consisted of four training trials (T), followed by a retention trial (RT) administered after a 30-minute rest. The test is repeated for five consecutive days. The number of errors in finding the platform for the last three days of testing was averaged and used for statistical analysis. RAWM was conducted using C57BL/6, twenty-five days after infection.

### RNAseq of Eco-HIV-infected mice brains

Upon sacrifice, brain tissue was collected from the mouse brain and frozen immediately in liquid nitrogen. Total RNA from fresh frozen brain samples was sent to Novogene Co. (Durham, NC) for next-generation sequencing (mRNA sequencing). All detailed information is described on their website: https://www.novogene.com. Briefly, the workflow included initial PolyA selection-based mRNA enrichment, mRNA fragmentation followed by random priming with subsequent first-and second-strand complementary DNA (cDNA) synthesis, library preparation, and sequencing using an Illumina NovaSeq 6000 sequencing system and 150 bp paired-end reads. The resulting data was checked for its quality, aligned to the mouse reference genome GRCm39, and differentially expressed genes were calculated by Novogene using the limma package (Bioconductor) in R.

### Microarray analysis of Eco-HIV-infected brains

Microarray experiments were conducted as previously described.^85^ Briefly, the microarray hybridization was performed at the Bionomics Research and Technology Center in EOSHI University of Medicine and Dentistry of New Jersey. RNA qualities were assessed by electrophoresis using the Agilent Bioanalyzer 2100 and spectrophotometric analysis before cDNA synthesis. Fifty nanograms of total RNA from each sample were used to generate a high-fidelity cDNA for array hybridization using NuGen WT-Ovation Pico RNA Amplification. After fragmentation and biotin labeling using NuGen Encore Biotin Module, the samples were hybridized to Affymetrix Mouse Gene ST Array Plate (Affymetrix, Santa Clara, CA). Washing and staining of all arrays were carried out in the Affymetrix fluidics module as per the manufacturer’s protocol. The detection and quantification of target hybridization was performed with an Affymetrix GeneChip Scanner. Data analysis was performed as described^85^. The .cel data files generated by the Affymetrix microarray hybridization platform were analyzed by the GeneSpring GX software (Agilent Technologies, Santa Clara, CA). Probe-level analysis was performed using the Robust Microarray Average (RMA) algorithm. Genes showing unpaired t-test p-values of <0.05 were defined as significantly changed.

### Glycomic analyses of mouse brains

A piece of flash-frozen brain was washed three times with PBS by adding 500µl PBS and centrifuged at 800xg for 5 minutes at 4°C. The piece was then homogenized using a cell homogenizer in PBS with protease inhibitor cocktail (Thermo Fisher; Catalog# 87786) and followed by centrifugation at 4°C for 5 minutes at 1500xg. After discarding supernatant, 300µl of PBS containing TritonX and protease inhibitor cocktail was added to the pellet and sonicated for 1 minute at high intensity. Cell lysate was then centrifuged for 10 minutes at 12000xg at 4°C and the supernatant containing protein was recovered and quantified using micro-BCA kit (Thermo Fisher; Catalog# 23235). Samples were normalized based on protein amount, and 0.4μg of protein was labeled with Cy3 (Cytivia; Catalog# PA23001). Excess of Cy3 was removed using desalting columns (Thermo Fisher; Catalog# 89883). Labeled proteins were hybridized overnight to lectin microarrays containing 90 lectins. The resulting chips were scanned for fluorescence intensity on each lectin-coated spot using a fluorescence scanner Bio-Rex Scan 200, and data were normalized using the global normalization method.

### Measurement of inflammatory markers in the brain and blood of mice infected or not with Eco-HIV

The concentration of 43 cytokines in mouse plasma and brain lysate was determined using the OLINK Target 48 Mouse Cytokine panel (OLINK; Catalog# 93400). Mouse brain tissue was thawed in 5% *w/v* T-PER Tissue Protein Extraction Reagent (ThermoScientific; Catalog# 78510) containing 1% protease/phosphatase inhibitor cocktail (Sigma-Aldrich; Catalog# 11697498001) and homogenized using a bead-based TissueLyzer II (QIAGEN) at 50Hz for 5 min. Samples were subsequently centrifuged at 10,000g for 10 min at 4°C to remove cell/tissue debris and soluble protein fraction was collected. Brain lysate protein concentrations were determined using a Pierce bicinchoninic acid assay (ThermoScientific; Catalog# 23225), adjusted to 1000μg/ml, and stored at -80°C until use. Thawed plasma and brain lysates were incubated with DNA barcode-tagged antibody pairs. Amplification and quantitation of antigen-specific DNA barcodes were then carried out by multiplex PCR on an OLINK Signature Q100, and calibration was used to determine absolute cytokine concentration.

### Assessment of inflammation markers on EVs

EVs were isolated from the unfixed frozen brain tissues as previously.^86^ To isolate EVs, frozen tissues were briefly sliced on dry ice and then dissociated in 3 ml Hibernate-E solution (Gibco, Catalog# A1247601) containing 20 units Papain (Worthington, catalog# LK003178) in Earle’s Balanced Salt Solution (Gibco; Catalog# 14155063) at 37°C for 15 min. After incubation, the tissue samples were immediately added with ice-cold Hibernate E with Halt Protease and Phosphatase Inhibitor cocktail (Catalog# PI178443) to stop digestion. Homogenized tissues were filtered through a sterile 40-μm mesh filter (Thermo Fisher; Catalog# 22-363-547) and sequentially centrifuged at 300 xg for 10 min at 4°C. Supernatants were transferred to fresh tubes and centrifuged at 2000 xg for 15 min at 4°C to remove apoptotic bodies. The resulting supernatants were then centrifuged at 13,000 x g for 30 min at 4°C, and the obtained EVs were stored at -80°C until flow cytometry analysis (**Supplementary Fig. 6a**).

EV quantity and surface markers were measured after samples were thawed and subjected to a freeze–thaw cycle. EVs were stained with pre-titrated volumes of fluorochrome-conjugated monoclonal antibodies: neuronal proteins Tau-Alexa Fluor 647 (Tau 46, Santa Cruz, Catalog# sc-32274 AF647) and β-Amyloid-PE (Santa Cruz, B-4, Catalog# sc-28365 PE), and CCR5-PE/Cy7 (clone: HM-CCR5, BioLegend, Catalog# 107018). Antibodies were filtered through a 0.22 µm centrifugal filter (Millipore) to remove aggregates. Fluorescence minus one (FMO) controls were included to determine the background signal. One to 2 µL of each antibody were added to 20 µL of EVs and incubated at 4°C for 30 minutes. EVs were then diluted in 0.22 µm-filtered PBS to appropriate dilutions to avoid coincident detection, as previously described.^87^

In accordance with the methodological guidelines for EV studies,^88^ EVs were characterized using a high-sensitivity Aurora spectral flow cytometer equipped with five lasers and enhanced small particle detection module (Cytek Biosciences). The MIFlowCyt checklist was included in **Supplementary Table 1**. Quality control was performed using SpectroFlo QC Beads according to the manufacturer’s instructions (Cytek Biosciences). A clean flow cell procedure was conducted prior to sample analysis to minimize EV carryover. A 0.22 µm-filtered PBS control was recorded to estimate the background noise. Unstained samples were used under the same conditions as the stained EV samples to ensure accurate unmixing and quantification of autofluorescence by flow cytometry. Side scatter was measured using the 405nm violet laser at a threshold of 1000 arbitrary units. Samples were acquired for 60 seconds at a low flow rate (∼15 µL/min). The reference bead mix (Apogee Flow Systems, Catalog# 1527) containing defined sizes of polystyrene (80, 110, 500 nm) and silica (180–1300 nm) beads was used to evaluate fluorescence performance and establish EV gates, as previously published.^89,90^ Representative flow cytometry plots showing the expression of surface markers are shown in **Supplementary Fig. 6b**. EV counts/µL were calculated using the flow rate of the cytometer. Analysis was performed using SpectroFlo software (Cytek Biosciences).

### Statistical analyses

For longitudinal data in Fig. 1, biomarkers with >40% unquantifiable values were dichotomized; those with 40-15% unquantifiable values were transformed to quartiles, and those with high skew and/or kurtosis were rank-transformed to deciles. Normally distributed biomarkers were standardized before modeling. Mixed models with fixed and random effects (for intercepts and time) were fit with CI (impaired vs. unimpaired) and continuous NPZ4 as exposures. Continuous outcomes were analyzed with linear regression, decile-and quartile-transformed outcomes were analyzed with ordinal logistic regression, and dichotomized biomarkers were analyzed with Robust Poisson regression and a log link to estimate risk ratios. Analyses were performed in SAS version 9.4 (SAS Institute). For Supplementary Fig. 2, linear models were fit in SAS. Analyses for Figs 2–6 were performed in GraphPad Prism (version 10). The specific analysis used for each figure is described in the corresponding legend.

### Data availability

All data are available within the manuscript and its figures and/or upon request.

## SUPPLEMENTARY FIGURE LEGENDS

**Supplementary Figure 1.** Capillary electrophoresis analysis of IgG and plasma *N*-glycans from females and males living with HIV on ART. Representative electropherogram showing resolved glycan peaks and their corresponding annotated structures for (a) IgG and (b) plasma *N*-glycans. Each peak corresponds to a distinct glycan species identified based on migration time.

**Supplementary Figure 2.** Validation of HIV-CI–associated glycomic alterations in an independent cross-sectional subset of the ACTG cohort HAILO. (a) Linear model results showing associations between glycans and CI status. Red = glycans positively associated with CI, blue = glycans associated with no-CI status. (b) Linear model results showing associations between glycans and NPZ4 scores. Red = glycans correlated with lower NPZ4 scores, blue = glycans correlated with higher NPZ4 scores

**Supplementary Figure 3.** Representative gating strategy for measuring T cell activation and exhaustion markers in humanized mice. Live cells were first gated, followed by the identification of human CD45 CD3 leukocytes. These were further subdivided into CD4 and CD8 T cell subsets. Within the CD8 population, expression of activation markers (CD38, HLA-DR, CD69) and exhaustion markers (PD-1) was quantified, with isotype controls used to define gating boundaries.

**Supplementary Figure 4.** EcoHIV infection alters glycosylation-related gene expression in the mouse brain. 129×1/SvJ mice were inoculated intraperitoneally with EcoHIV or PBS (controls). One month post-infection, mice were sacrificed, and brain RNA was analyzed using gene microarray. Relative expression levels of NEU1 (encoding sialidase/neuraminidase) and multiple sialyltransferases were assessed. Heatmap presentation shows differential expression, with red indicating higher expression and green indicating lower expression. EcoHIV infection was associated with significantly increased NEU1 expression and reduced expression of several sialyltransferases, consistent with enhanced glycan degradation and impaired glycan synthesis. Statistical significance was determined using unpaired t-tests.

**Supplementary Figure 5.** Sialidase inhibitors prevent Eco-HIV–mediated systemic inflammation. A heatmap of plasma samples showing protein levels of several inflammatory markers in Eco-HIV–infected and control mice, with or without sialidase inhibitor treatment. Eco-HIV infection upregulated several of these markers, while sialidase inhibitors prevented these elevations. Red = higher expression; blue = lower expression. Mann–Whitney T tests.

**Supplementary Figure 6.** Measuring single EVs in brain tissues. **(a)** Schematic of the method for EV isolation and characterization. (b) Representative flow plots showing EV subtypes according to their surface marker expression.

**Supplementary Table 1.** MIFlowCyt checklist for EV experiments

## AUTHOR CONTRIBUTIONS

M.A-M conceived the study. D.V., L.C.N and M.A-M acquired the funding. L.B.G, A.B, E.H, E.G.M.M, T.A.P, H.H, S.T.Y, S.S, and C.F carried out experiments. J.G, E.B, D.D, M.B, A.S, N.S, and P.W.D performed and interpreted the humanized mouse experiments. H.T provided the lectin microarray. F.P, L.C.N, P.J.N, and K.T selected clinical samples and interpreted data. D.J.V supervised the Eco-HIV experiments. M.J.C performed the epigenetic clock analyses. J.G analyzed the human cohort data. L.B.G and M.A-M wrote the manuscript, and all authors reviewed and edited the final version.

## ACKNOWLEDGMENTS

We would like to thank study participants. This study is supported by the National Institutes of Health (NIH) R01NS117458 grant to M.A-M, L.C.N, and D.V as well as R01AG092241 and R01AI189353 to M.A-M. The study was also supported by NWCS 539 from ACTG to L.G.B. M.A-M is also supported by NIH grants (R01AI165079, R01AA029859, and R01DK123733). M.A-M is a member of the NIH-funded BEAT-HIV Martin Delaney Collaboratory to cure HIV-1 infection (1UM1Al126620). L.C.N has research time supported by R01AG063846 and P.W.D. has research time supported by P20GM103427 and R15AI178516. Research reported in this publication was supported by the National Institute of Allergy and Infectious Diseases of the NIH under Award Number UM1 AI068634, UM1 AI068636, UM1 AI106701, and AI069494. The content is solely the responsibility of the authors and does not necessarily represent the official views of the National Institutes of Health. We would like to thank the ACTG Statistical and Data Management Center (SDMC) for performing part of the statistical analyses.

