## Supplementary figures and images for "Inhibiting Glycan Degradation Prevents HIV-Induced Inflammaging and Cognitive Impairment"

### Supplementary Figure 1

**Supplementary Figure 1**

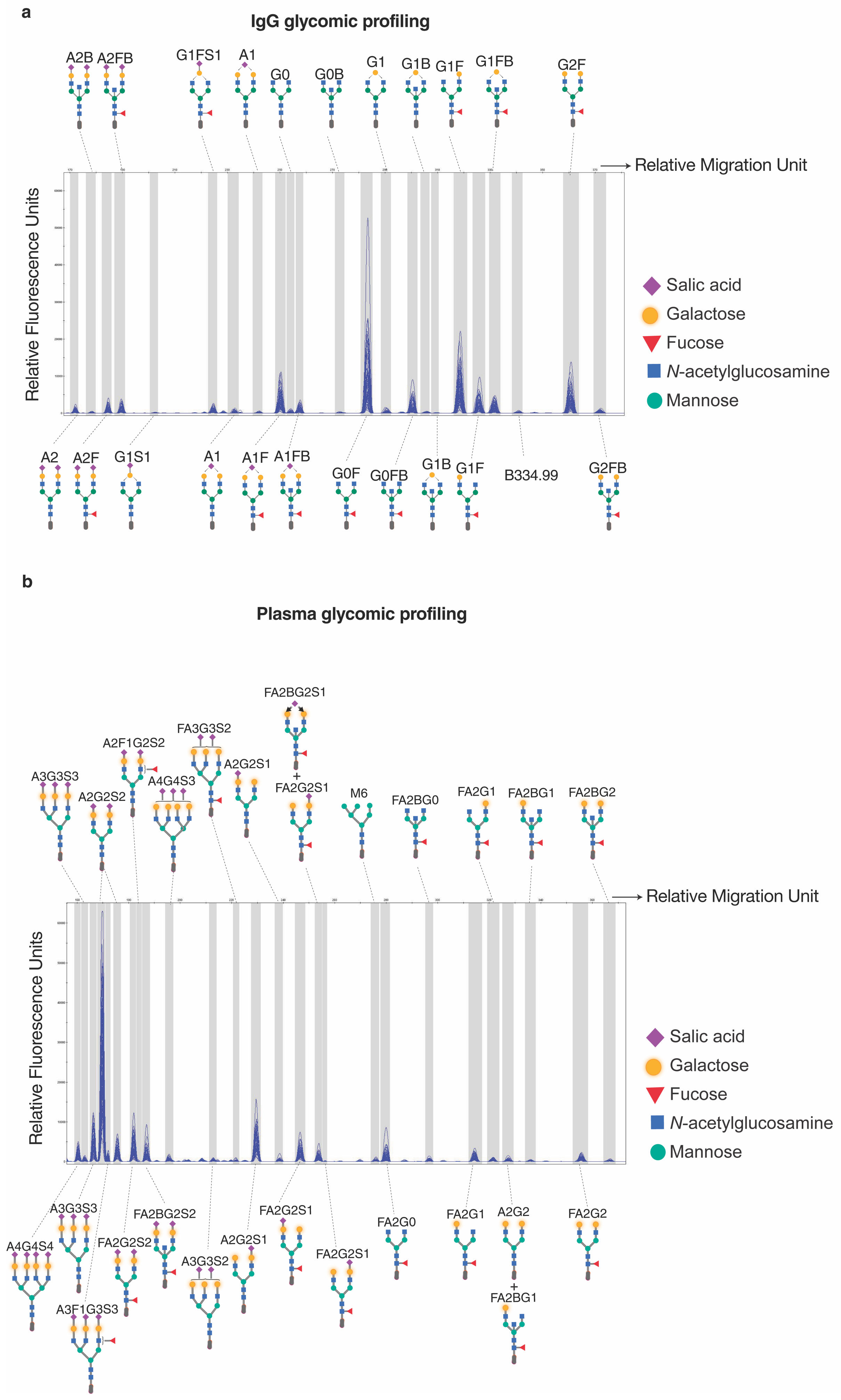

### Supplementary Figure 2

**Supplementary Figure 2**

**a**

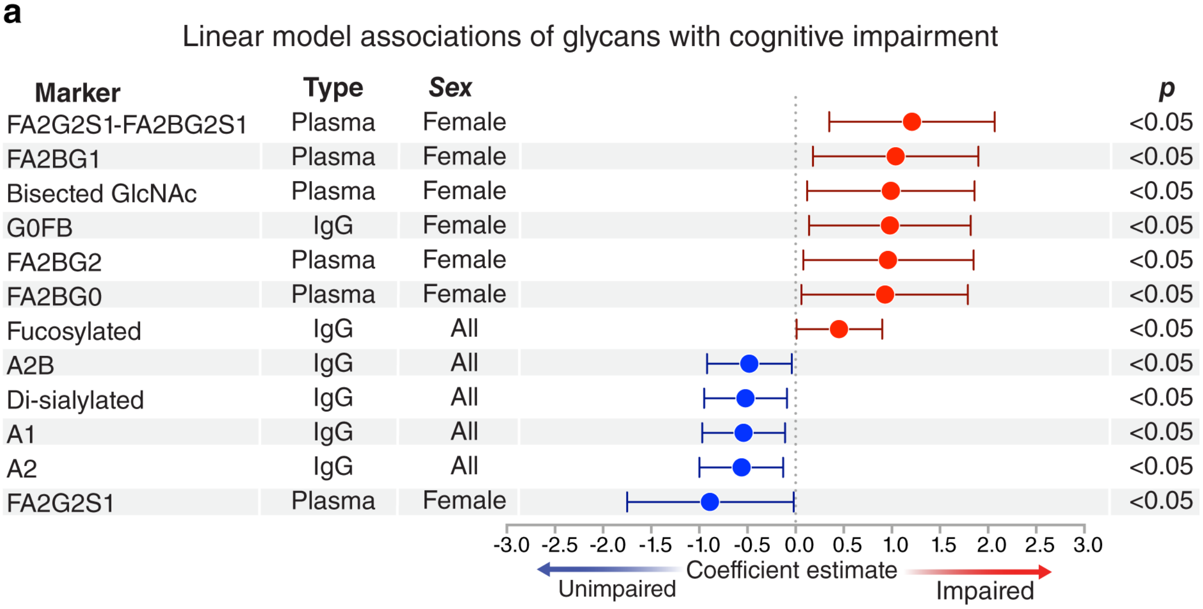


**b**

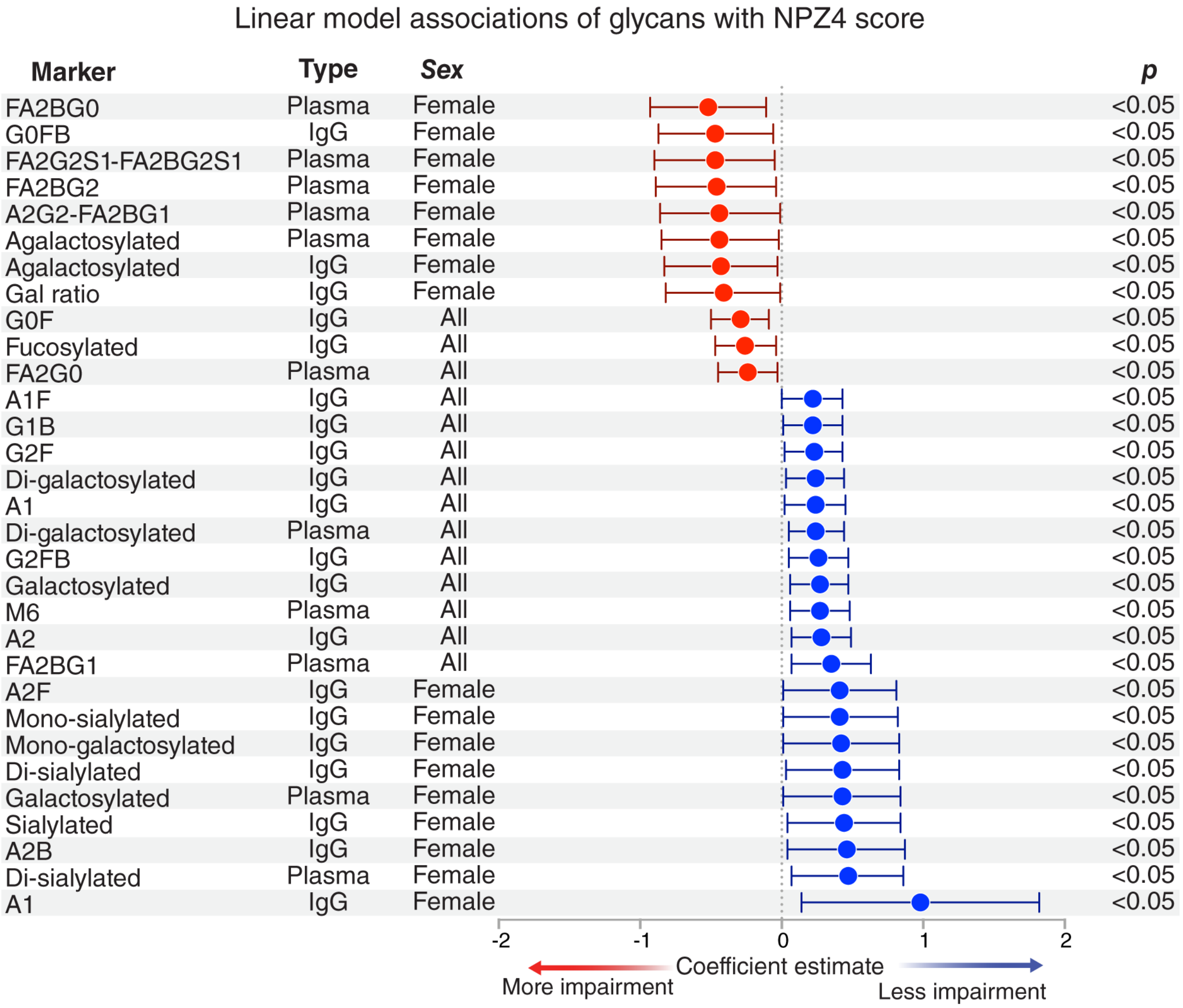

### Supplementary Figure 3

**Supplementary Figure 3**

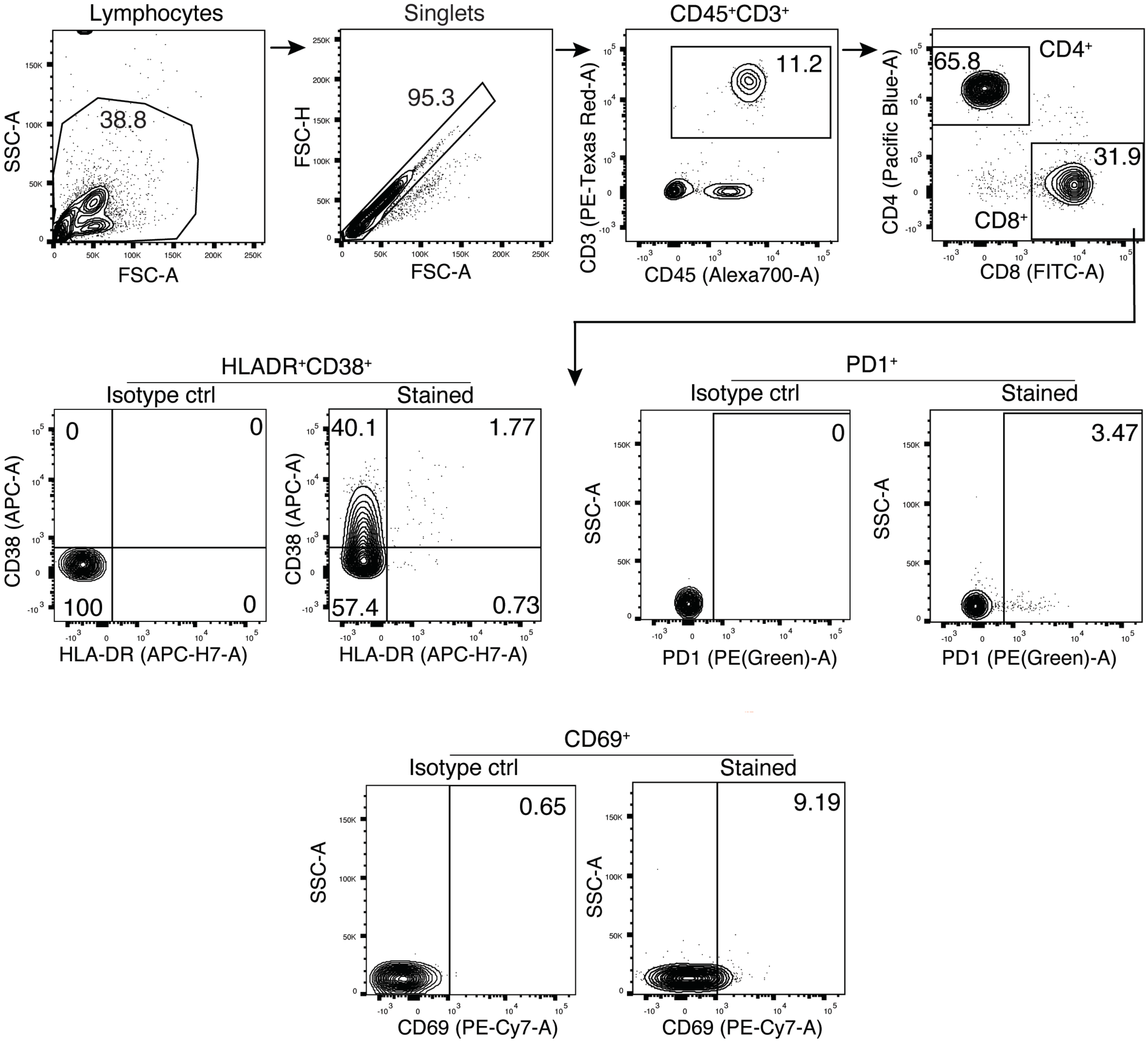

### Supplementary Figure 4

#### Supplementary Figure 4

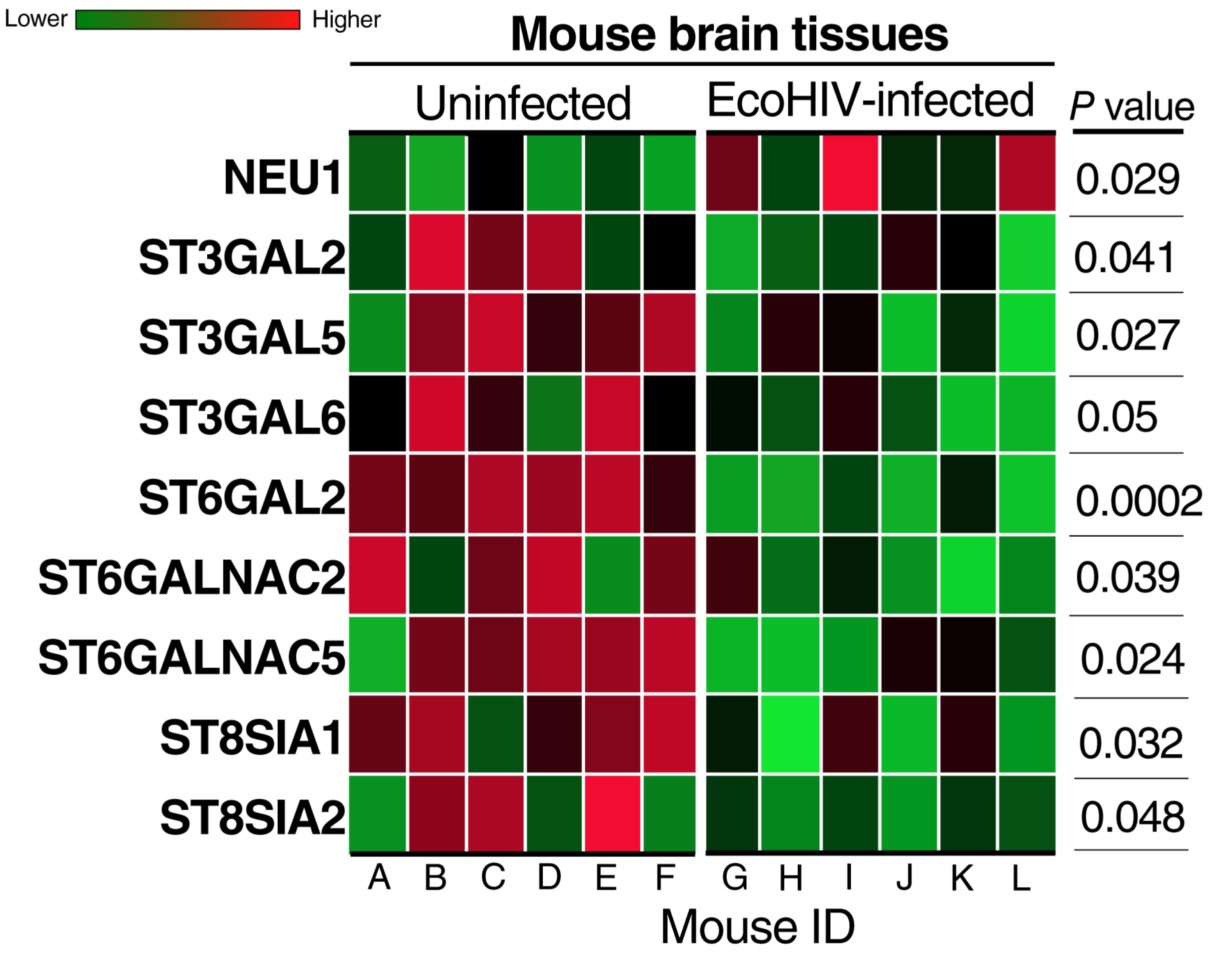

### Supplementary Figure 5

#### Supplementary Figure 5

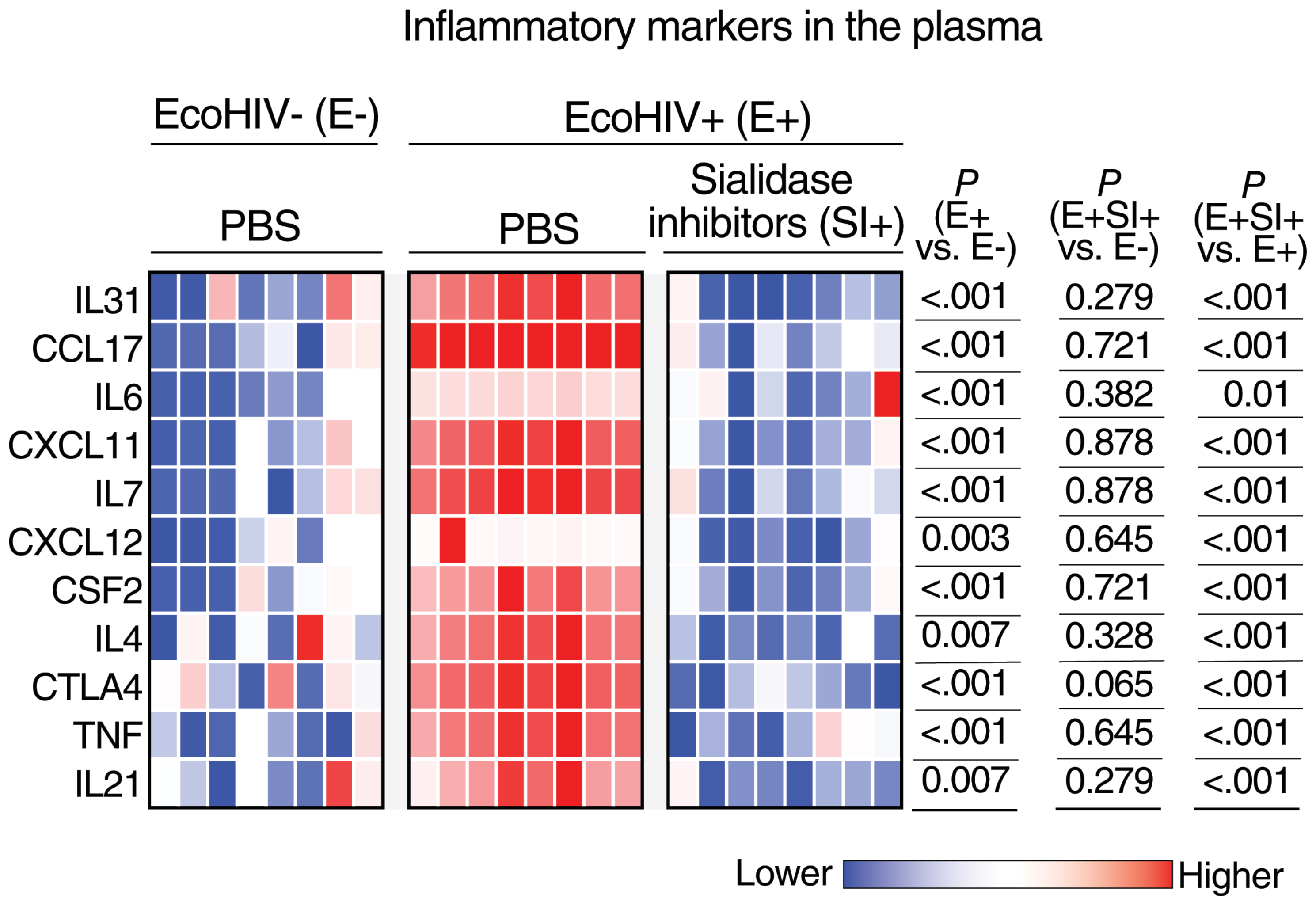

### Supplementary Figure 6

#### Supplementary Figure 6

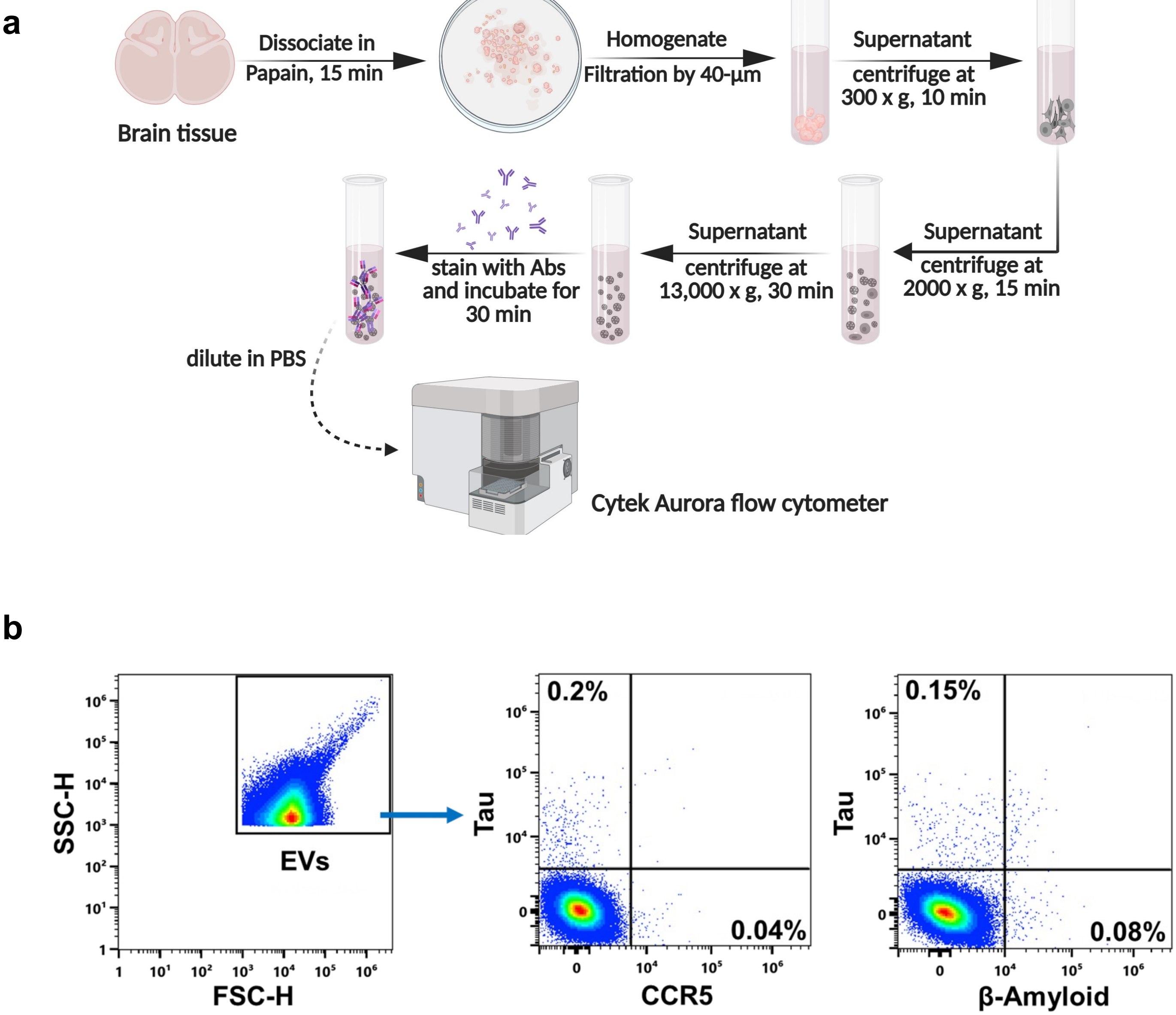
