## Supplementary Table 1 for "Inhibiting Glycan Degradation Prevents HIV-Induced Inflammaging and Cognitive Impairment"

**Supplementary Table 1.** MIFlowCyt checklist for EV experiments

| **Framework Criteria** | **What to report** | **Please complete each criterion** |
|---|---|---|
| 1.1 Preanalytical variables conforming to MISEV guidelines. | Preanalytical variables relating to EV sample including source, collection, isolation, storage, and any others relevant and available in the performed study. | Frozen brain tissues were dissociated in Hibernate-E with 20 U Papain (Earle's Balanced Salt Solution) and digestion stopped with Hibernate-E containing Halt Protease/Phosphatase inhibitor. EVs were isolated by stepwise centrifugation (300 × g, 10 min; 2000 x g, 15 min; 13,000 x g, 30 min; all at 4°C) and stored at -80°C until flow cytometry analysis. |
| 1.2 Experimental design according to MIFlowCyt guidelines. | EV-FC manuscripts should provide a brief description of the experimental aim, keywords, and variables for the performed FC experiment(s) using MIFlowCyt checklist criteria: 1.1, 1.2, and 1.3, respectively. Template found at www.evflowcytometry.org. | **1.1 Aim**: Measure EV quantity and putative cell source markers in brain tissues. **1.2 Keywords:** Extracellular vesicles, neuropathological markers. **1.3 Experimental variables:** EV subtypes were characterized in brain tissues from mice infected or not with Eco-HIV and treated or not with sialidase inhibitors using spectral flow cytometry. Scatter-based triggering was used for the detection of EVs. Sialidase inhibitor treatment was associated with reduced levels of brain-EVs expressing neurodegeneration-related markers. |
| 2.1 Sample staining details | State any steps relating to the staining of samples. Along with the method used for staining, provide relevant reagent descriptions as listed in MIFlowCyt guidelines (Section 2.4 Fluorescence Reagent(s) Descriptions). | EVs were stained with pre-titrated fluorochrome-conjugated antibodies: Tau-Alexa Fluor 647, β-amyloid-PE, and CCR5. Antibodies were filtered (0.22 μm) to remove aggregates. Fluorescence-minus-one controls were included. Antibodies (1-2 μL) were added to 20 μL EVs, incubated 30 min at 4°C, and diluted in 0.22-μm PBS to prevent coincident detection. |
| 2.2 Sample washing details | State any steps relating to the washing of samples. | Not applicable |
| 2.3 Sample dilution details | All methods and steps relating to sample dilution. | Brain-derived EVs were serially diluted (1:1, 1:10, 1:100, and 1:1000) in 0.22 μm-filtered PBS. |
| 3.1 Buffer alone controls. | State whether a buffer-only control was analyzed at the same settings and during the same experiment as the samples of interest. If utilized it is recommended that all samples be recorded for a consistent set period of time e.g. 5 minutes, rather than stopping analysis at a set recorded event count e.g. 100,000 events. This allows comparisons of total particle counts between controls and samples. | A 0.22-μm-filtered PBS-only control was analyzed under the same settings and during the same experiment as the study samples. The buffer-only control and all samples were acquired for 60 seconds at a low flow rate. |
| 3.2 Buffer with reagent controls. | State whether a buffer with reagent control was analyzed at the same settings, same concentrations, and during the same experiment as the samples of interest. If used state what the results were. | Buffer with reagents was recorded under the same acquisition settings, at the same concentrations, and during the same experiment as all study samples. No background signals. |
| 3.3 Unstained controls. | State whether unstained control samples were analyzed at the same settings and during the same experiment as stained samples. If used, state what the results were, preferably in standard units. | Unstained controls were measured at the same dilution as matched stained samples. |
| 3.4 Isotype controls. | The use of isotype controls is applicable to immunofluorescence labelling only. State whether isotype controls were analyzed at the same settings and during the same experiment as stained samples. If utilized, state which antibody they are matched to, the concentration used, and what the results were (Section 4.2, 4.3, 4.4). Due to conjugation differences between manufacturers if should be stated if the isotype controls are from the same manufacturer as the matched antibodies. | Isotype controls were not measured. |
| 3.5 Single-stained controls. | State whether single-stained controls were included. If used state whether the single-stained controls were recorded using the same settings, dilutions, and during the same experiment as stained samples and state what the results were, preferably in standard units (Section 4.2, 4.3, 4.4). | Single stain controls were included and used as reference controls for the unmixing. |
| 3.6 Procedural controls. | State whether procedural controls were included. If used, state the procedure and if the procedural controls were acquired at the same settings and during the same experiment as stained samples. | Procedural controls (buffer only, buffer with reagents, unstained samples, and fluorescence-minus-one controls) were acquired under the same settings and in the same experiment as stained samples. |
| 3.7 Serial dilutions. | State whether serial dilutions were performed on samples and note the dilution range and manner of testing. The fluorescence and/or scatter signal intensity would ideally be reported in standard units (see Section 4.3, 4.4) but arbitrary units can also be used. This data is best reported by plotting the recorded number events/concentration over a set period of time at different sample dilution. The median fluorescence intensity at each of the dilutions should also ideally be plotted on the same or a separate plot. | Samples were serially diluted (1:1, 1:10, 1:100, and 1:1000) in 0.22-μm-filtered PBS and run for 60 seconds at low flow rate; the dilution yielding single EV detection (abort rate <10% of count rate) was used to avoid swarming. |
| 3.8. Detergent treated EV-samples | State whether samples were detergent treated to assess lability. If utilized, state what detergent was used, the end concentration of the detergent, and what the results were of the lysis. | Study samples were not treated with detergent. |
| 4.1 Trigger Channel(s) and Threshold(s). | The trigger channel(s) and threshold(s) used for event detection. Preferably, the fluorescence calibration (Section 4.3) and/or scatter calibration (Section 4.4) should be used in order to report the trigger channel(s) and threshold(s) in standardized units. | 405nm violet laser; threshold of 1000 arbitrary units. |
| 4.2 Flow Rate / Volumetric quantification. | State if the flow rate was quantified/validated and if so, report the result and how they were obtained. | Samples were acquired at the lowest flow rate (15 μL/min) as measured using the internal flow rate sensor of the cytometer. |
| 4.3 Fluorescence Calibration. | State whether fluorescence calibration was implemented, and if so, report the materials and methods used, catalogue numbers, lot numbers, and supplied reference units for the standards. Fluorescence parameters may be reported in standardized units of MESF, ERF, or ABC beads. The type of regression used, and the resulting scatter plot of arbitrary data vs standard data for the reference particles should be supplied. | Calibration was performed using SpectroFlo QC Beads (catalog number: SKU B7-10001, lot: 2006, Cytek Biosciences). |
| 4.4 Light Scatter Calibration. | State whether and how light scatter calibration was implemented. Light scatter parameters may be reported in standardized units of nm2, along with information required to reproduce the model. | A reference bead mix (cat. no. 1493, Apogee Flow Systems) and NIST-traceable polystyrene beads (3000 Series Nanosphere Size Standards, Thermo Fisher Scientific) were used for calibration. Data were analyzed with SpectroFlo software (v2.2.0.4; Cytek). |
| 5.1 EV diameter/surface area/volume approximation. | State whether and how EV diameter, surface area, and/or volume has been calculated using FC measurements. | NA |
| 5.2 EV refractive index approximation. | State whether the EV refractive index has been approximated and how this was done. | NA |
| 5.3 EV epitope number approximation. | State whether EV epitope number has been approximated, and if so, how it was approximated. | NA |
| 6.1 Completion of MIFlowCyt checklist. | Complete MIFlowCyt checklist criteria 1 to 4 using the MIFlowCyt guidelines. Template found at www.evflowcytometry.org. | The MIFlowCyt checklist was attached in the Supplementary Information. |
| 6.2 Calibrated channel detection range | If fluorescence or scatter calibration has been carried out, authors should state whether the upper and lower limits of a calibrated detection channel were calculated in standardized units. This can be done by converting the arbitrary unit scale to a calibrated scaled, as discussed in Section 4.3 and 4.4, and providing the highest unit on this scale and the lowest detectable unit above the unstained population. The lowest unit at which a population is deemed 'positive' can be determined a variety of ways, including reporting the 99th percentile measurement unit of the unstained population for fluorescence. The chosen method for determining at what unit an event was deemed positive should be clearly outlined. | NA |
| 6.3 EV number/concentration. | State whether EV number/concentration has been reported. If calculated, it is preferable to report EV number/concentration in a standardized manner, stating the number/concentration between a set detection range. | EV number was reported in the Results section. |
| 6.4 EV brightness. | When applicable, state the method by which the brightness of EVs is reported in standardized units of scatter and/or fluorescence. | NA |
| 7.1. Sharing of data to a public repository. | Provide a link to the experimental data in a public data repository. | NA |
